# Astrocytes adopt a progenitor-like migratory strategy for regeneration in adult brain

**DOI:** 10.1101/2024.05.18.594292

**Authors:** Marina Herwerth, Matthias T. Wyss, Nicola B. Schmid, Jacqueline Condrau, Luca Ravotto, José María Mateos Melero, Andres Kaech, Gustav Bredell, Carolina Thomas, Christine Stadelmann, Thomas Misgeld, Jeffrey L. Bennett, Aiman S. Saab, Sebastian Jessberger, Bruno Weber

## Abstract

Mature astrocytes become activated upon non-specific tissue damage and contribute to glial scar formation. Proliferation and migration of adult reactive astrocytes after injury is considered very limited. However, the regenerative behavior of individual astrocytes following selective astroglial loss, as seen in astrocytopathies, such as neuromyelitis optica spectrum disorder, remains unexplored. Here, we performed longitudinal *in vivo* imaging of cortical astrocytes after focal astrocyte ablation in mice. We discovered that perilesional astrocytes develop a remarkable plasticity for efficient lesion repopulation. A subset of mature astrocytes transforms into reactive progenitor-like (REPL) astrocytes that not only undergo multiple asymmetric divisions but also remain in a multinucleated interstage. This regenerative response facilitates efficient migration of newly formed daughter cell nuclei towards unoccupied astrocyte territories. Our findings define the cellular principles of astrocyte plasticity upon focal lesion, unravelling the REPL phenotype as a fundamental regenerative strategy of mature astrocytes to restore astrocytic networks in the adult mammalian brain. Promoting this regenerative phenotype bears therapeutic potential for neurological conditions involving glial dysfunction.

## Introduction

In response to various pathological triggers, astrocytes undergo a process called reactive astrogliosis. This action encompasses morphologic, molecular and functional changes^1–3^ and has been studied in detail in traumatic central nervous system (CNS) injury models, where reactive astrocytes remain stationary at the rim of lesion borders, proliferate and form a glial scar.^4–6^ Similar behavior has been found in other neurological conditions,^7^ such as stroke^8^, Alzheimer’s disease^9^, and neuroinflammation.^10^ Although this reaction is heterogeneous, the key features across different stimuli include cellular hypertrophy, upregulation of glial fibrillary acidic protein (GFAP), limited proliferation and activation of different transcription factors.^1^ Our knowledge about reactive astrocytes in the adult brain is mainly based on studies involving direct mechanical or hypoxic and global tissue damage. Under these conditions, a subset of mature astrocytes at the lesion border proliferates symmetrically for one cycle, rarely twice after repetitive traumatic brain injury, and upregulates some neural precursor cell (NPC) hallmark proteins.^11^ These adult-born daughter cells remain stationary in close vicinity to each other and stay within the astrocyte lineage.^4,11^ In brain pathologies involving vessel disrupture, these cells were restricted to the juxtavascular niche^4^, suggesting that blood-brain barrier (BBB) leakage is a trigger for astrocyte proliferation.^12^ Glial scar itself is a dysfunctional structure and its contribution to regeneration is still under debate.^6,12–14^ However, models to investigate responses to a selective astrocyte depletion without glial scar formation are lacking and the full regenerative capacity of mature astrocytes is unknown.

Here, we applied a new antibody-mediated strategy to ablate astrocytes in a selective and locally controlled manner in the intact adult brain. We targeted the water channel aquaporin-4 (AQP4), which is almost exclusively expressed by astrocytes in the CNS.^15^ Autoantibodies against AQP4 can be found in patients suffering from Neuromyelitis optica spectrum disorder (NMOSD) – a paradigmatic autoimmune disease of the CNS with B cell orchestrated immune responses.^16,17^ By tracking perilesional astrocytes with chronic, intravital two-photon microscopy in anesthetized and awake mice, we discover that mature cortical astrocytes adopt progenitor cell-like behaviors, including migration of adult-born daughter cell nuclei to unoccupied astrocytic territories. This self-renewal program is neither associated with the vascular niche, nor dependent on connexins 30/43 or NLPR3 inflammasome signaling. Applying an alternative all-optical apoptotic ablation method to astrocytes,^18^ we identify the astrocyte loss volume as a key driver of these regenerative mechanisms. These astrocytic features are also apparent in human AQP4-IgG-mediated lesions from NMOSD patients. We propose that in response to local astrocyte loss, regardless of the underlying cause, mature astrocytes transform into reactive progenitor-like (REPL) cells, which exhibit a fundamental regenerative competence for restoring astrocyte tiling.

### Astrocytes regenerate rapidly after local depletion

To assess astrocytic behavior over time after selective astrocyte depletion we used transgenic mice expressing the enhanced green fluorescent protein (EGFP) under the control of the pan-astroglial Aldh1l1 promoter.^19^ We have previously shown by acute *ex vivo* and *in vivo* two-photon imaging, that human derived recombinant AQP4-IgG efficiently binds and depletes mouse astrocytes within hours.^20,21^ Similarly, step-wise intracortical microinjection of AQP4-IgG and complement in Aldh1l1^GFP^ mice (Fig. 1a) led to cylindrical lesions devoid of astrocytes (Fig.1b) in the somatosensory cortex. Such lesions were absent after injection of complement with isotype control recombinant IgG (Ctrl-IgG; Fig.1b). The extent of astrocyte depletion was dependent on the antibody titer and the concentration of complement (Extended Data Fig. 1). This selective astrocyte depletion did not affect the morphology and function of the vasculature as visualized after intravenous administration of Texas Red dextran (Fig. 1b-c). No overt changes in vessel density, average vessel diameter (Fig. 1d-e), resting capillary blood flow, and capillary diameter measured by two-photon line scan imaging (Fig. 1f-g) were observed compared to Ctrl-IgG injections. Furthermore, immunohistological staining for the neuronal marker NeuN showed the presence of neuronal population within the astrocyte-depleted area (Extended Data Fig. 2).

**Fig. 1.**
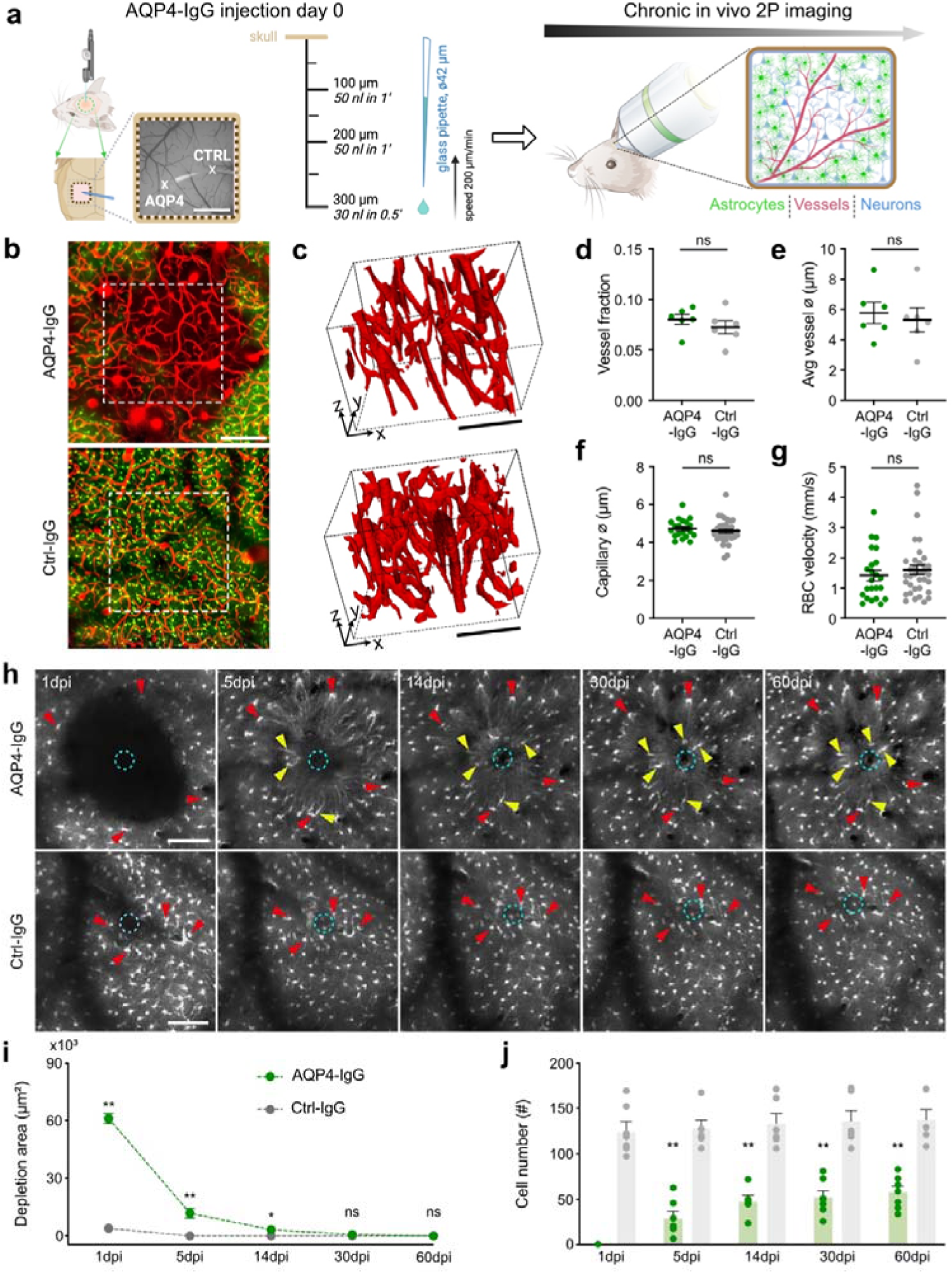
AQP4-IgG-based selective astrocyte depletion leads to efficient astrocytic lesion repopulation. **a,** Protocol of AQP4-IgG-based astrocyte depletion in the somatosensory cortex of anesthetized mice. Implantation of a permanent window allows chronic *in vivo* two-photon microscopy imaging (scale bar = 1 mm).**b**, Projected two-photon image z-stacks (100 µm) from Aldh1l1^GFP^ mice, demonstrating focal loss of astrocyte-expressing GFP signal (scale bar = 100 µm). Vessels were visualized with i.v. injection of Texas Red dextran (70kD). White dashed squares mark the locations of vascular segmentations shown in **c**. **c,** Vascular segmentations based on *in vivo* data from **b** (scale bar = 100 µm). **d-g,** AQP4-IgG-mediated ablation of astrocytes does not alter vascular structure compared to the intracortical injection of a control antibody (Ctrl-IgG). Quantification of vessel fractions in **d** and mean vessel diameters in **e** determined in the segmentation, n = 6 animals per condition (Mann-Whitney test) and average capillary diameters in **f** and red blood cell (RBC) velocities in capillaries in **g** measured *in vivo*, n = 23-35 vessels from 3 mice, respectively (Mann-Whitney test). **h,** Using longitudinal *in vivo* two-photon imaging in Aldh1l1^GFP^ mice, re-coverage of the astrocyte depleted area after AQP4-IgG-injection was followed over 60 days (upper panels). Cells in the direct vicinity of the lesion (perilesional astrocytes, red arrowheads) remain in their location throughout the experimental time period. Within the initially astrocyte-depleted area, an increasing number of newly appearing astrocytes are detected (yellow arrowheads) over time. Lower panels: After Ctrl-IgG injection, cells in the direct vicinity of the injection site (cyan dashed circles) are present throughout the whole follow-up period (red arrowheads, scale bars = 100 µm). **i-j,** Quantification of mean depletion area size over time in **i** and the cell number in the lesion area in **j**. n = 6-8 areas from 6-7 animals per group, *p < 0.05, **p < 0.01, and not significant (ns; Mann-Whitney test). Data are represented as mean ± SEM.

We next tracked astrocytic responses after local astrocyte depletion via longitudinal *in vivo* two-photon microscopy and found that the area of astrocyte GFP signal loss was completely restored within a few weeks (Fig. 1h-i). In parallel, the number of astrocytic cell bodies in the depleted area rapidly increased, reaching a plateau within a few weeks (Fig. 1j). Importantly, focal ablation of astrocytes did not cause the formation of a glial scar, as seen in animal models of traumatic brain injury or stroke.^4,5,8^

**Extended Data Fig. 1.**
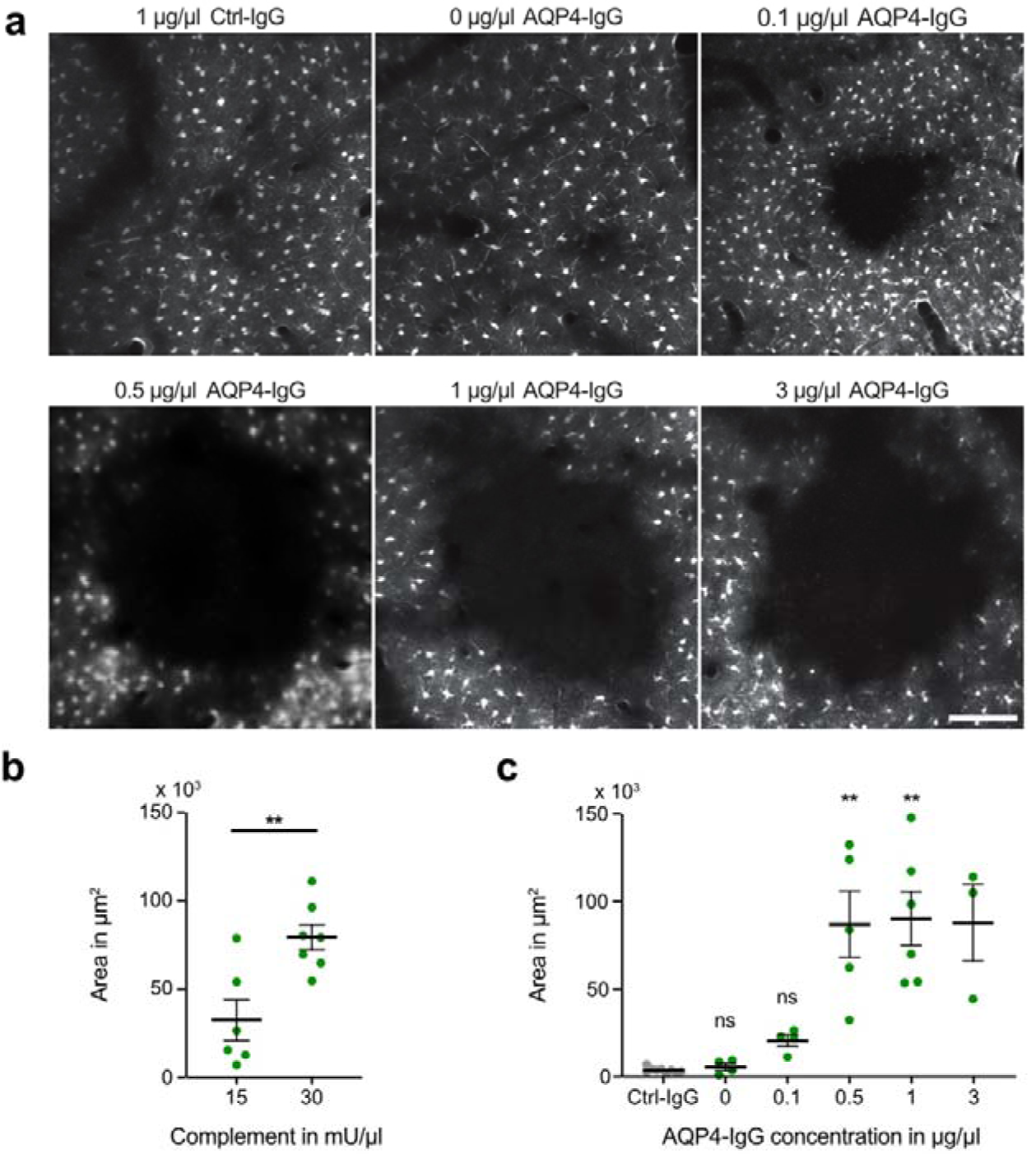
Titration of AQP4-IgG and complement for lesion induction. **a,** Examples of astrocyte depletion areas 1 day after injection of different AQP4-IgG concentrations (complement 30 mU/µl; scale bar = 100 µm). **b,** Depletion areas with two different complement concentrations (and a fixed concentration of 1µg/µl AQP4-IgG), n = 4-5 animals per group. **p < 0.01 (Mann-Whitney test). **c,** Astrocyte depletion areas with different AQP4-IgG concentrations and a fixed complement concentration (30 mU/µl), n = 3 – 9 animals per group. **p < 0.01 and ns = not significant (Kruskal-Wallis test). Data are represented as mean ± SEM.

**Extended Data Fig. 2.**
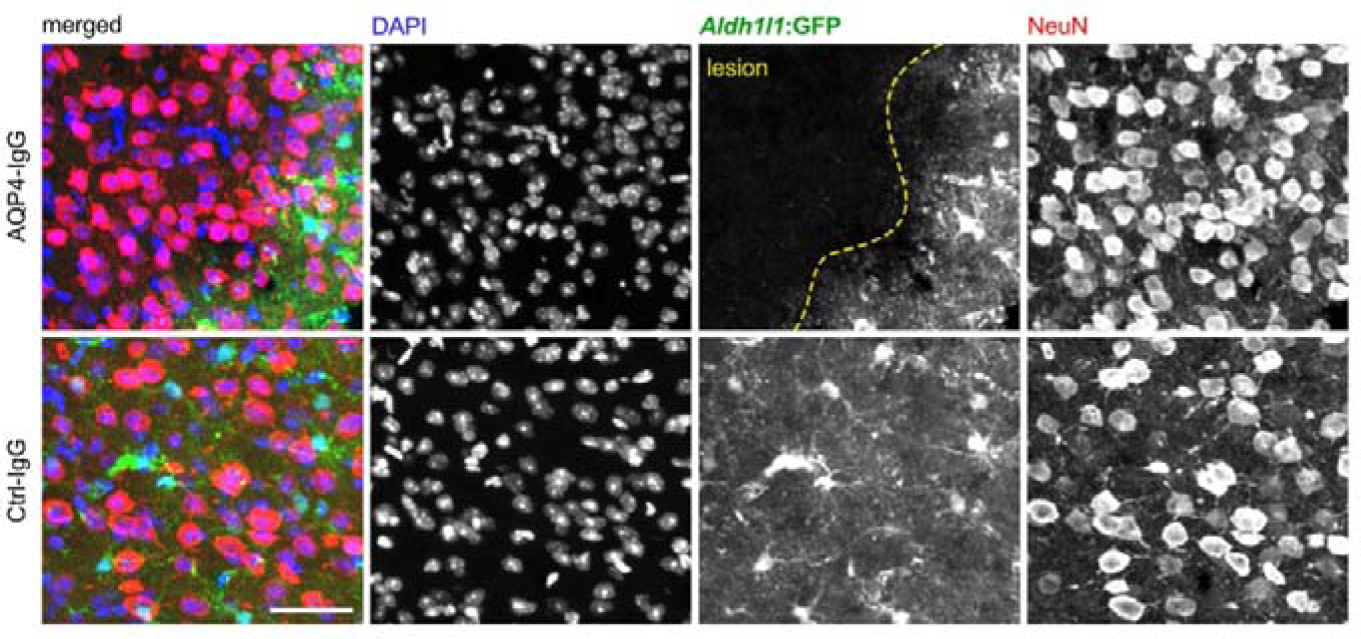
AQP4-IgG injection selectively depletes astrocytes. Confocal images of endogenous Aldh1l1^GFP^ signal and NeuN staining 1 day after AQP4-IgG (top) or Ctrl-IgG (bottom) injection reveal persistence of neurons in the lesion area (scale bar = 40 µm). The yellow dashed line indicates the lesion border.

### Early directed remodeling of perilesional astrocytes

Astrocytes show a reactive phenotype with heterogeneous morphological, transcriptional and functional changes on exposure to various pathological stimuli.^1^ To assess the dynamics of the reactivation of surviving perilesional astrocytes *in vivo*, we performed a 3D vector-based analysis of astrocyte process length and orientation of randomly chosen perilesional astrocytes (Fig. 2a-b). We found a robust elongation of astrocytic processes, reaching the maximum length at 5 days post injection (dpi) followed by a recovery at 60 dpi (Fig. 2c). This was accompanied by the specific orientation of astrocytic processes towards the center of the lesion (COL). In contrast to process elongation, this polarization remained stable within the two-month observation period (Fig. 2d). In line with these morphological changes, we found an upregulation of GFAP, a typical marker for astrogliosis,^22^ in histological sections comparing AQP4-IgG and Ctrl-IgG areas (Fig. 2e-f).

**Fig. 2.**
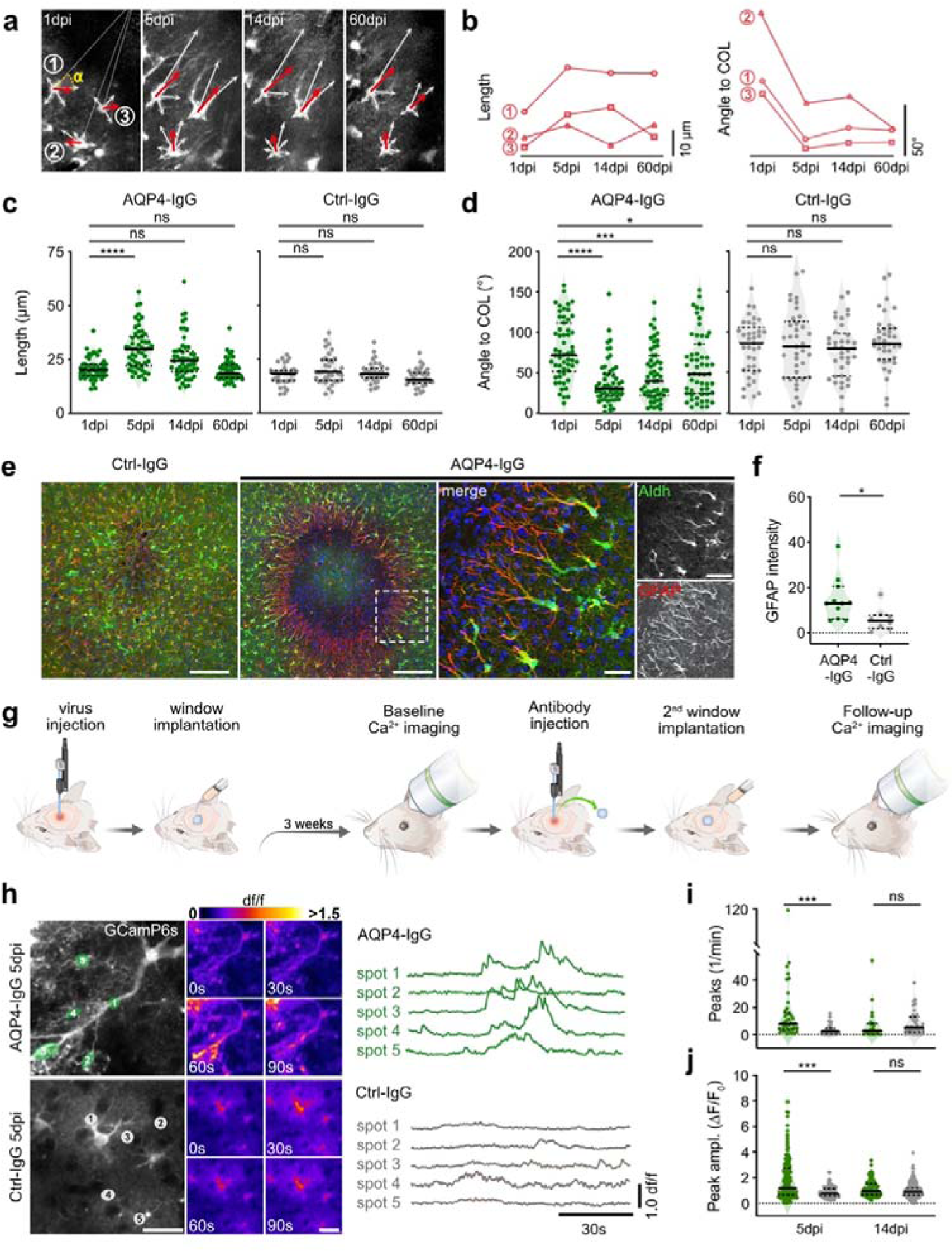
AQP4-IgG-mediated astrocyte reactivity shows early morphological and functional changes. **a-b,** Representative images and traces of three cells illustrating the vector-based analysis approach used for determination of the process length and angle IZ (yellow); dashed white lines project to the center of the lesion (COL). White arrows show individual vectors for main astrocytic processes and the respective mean vectors are indicated by red arrows. **c-d**, Vector-based quantification of astrocytic process lengths and polarization up to 60 dpi. n = 55 astrocytes from 3-4 lesions from 3 mice for AQP4-IgG, n = 37 astrocytes from 4 lesions from 4 mice for Ctrl-IgG, (Friedman test followed by Dunn’s multiple comparisons test). **e,** Confocal images of immunohistochemical staining against GFAP of fixed brain tissue 5 days after AQP4-IgG or Ctrl-IgG injection. Right panels: higher magnification images from the perilesion of the boxed area in the overview image, illustrating the upregulation of GFAP (DAPI in blue; scale bars = 100 µm in overview picture and 20 µm in zoom-ins). **f,** Corresponding quantification of GFAP area intensity, n = 8-11 mice per condition. (Mann-Whitney test). **g,** Schematic illustration of experimental workflow for imaging of astrocytic calcium activity in astrocytes before and after AQP4-IgG/Ctrl-IgG injection. **h,** Representative two-photon images of GCamP6s activity in astrocytes from an AQP4-IgG- and Ctrl-IgG-injected area (top and bottom respectively). Color-coded images display calcium levels in the same field of view at different time points (scale bars = 20 µm). On the right: individual traces from the indicated spots (1-5). **i-j,** Quantification of calcium peak frequency and maximal peak amplitudes at 5 and 14 dpi for AQP4-IgG compared to Ctrl-IgG, n = 6 – 7 mice at 5 dpi and 5 mice at 14 dpi, respectively (Kruskal-Wallis test). Thick black lines represent the median and fine black dashed lines represent the first and third quartiles. *p < 0.05, **p < 0.01, ***p < 0.01, ****p < 0.0001, and not significant (ns).

Astrocytes display distinct local calcium transients^23,24^ that may be dysregulated in pathological conditions.^25,26^ To examine whether perilesional astrocytes undergo functional alterations in their calcium signaling, we measured spontaneous astrocytic calcium transients during lesion recovery, using an AAV-based GCaMP6s injected 3 weeks before AQP4-IgG/Ctrl-IgG injection (Fig. 2g). While at 1 dpi, the spontaneous activity of perilesional astrocytes was similar in both conditions (data not shown), at 5 dpi, perilesional astrocytes developed a marked increase in calcium transient frequencies and amplitudes compared to astrocytes from Ctrl-IgG-injected areas (Fig. 2h-j, Movie 1). Two weeks later (at 14 dpi) when lesions were almost completely repopulated, calcium signaling normalized (Fig. 2i-j). This implies that the transformation of perilesional astrocytes is accompanied by transient changes in intracellular calcium dynamics.

### Perilesional astrocytes undergo multiple asymmetric cell divisions

Next, we wondered about the origin of newly appearing astrocytes in the lesions. Given that tracking single astrocytes with global Aldh1l1^GFP^ labeling is difficult, we sparsely labeled astrocytes with a low-dose tamoxifen injection in GLAST^CreERT2^ x ROSA26^tdTomato^ mice in combination with an intravenous injection of PHP.eB/2-hGFAP-eGFP viruses to achieve widespread astrocytic labeling in order to precisely localize the lesion borders (Fig. 3a). Five days after lesion induction, the volume occupied by a single astrocyte increased by ∼70% while the volume of astrocytes from control areas remained stable (Fig. 3b). Corroborating the observations made in Aldh1l1^GFP^ mice, this volume change was dominated by a thickening of main processes and increased complexity of branch morphology (Fig. 3b-d). By tracking individual reactive astrocytes over several weeks, we also observed focal swellings within the thick processes at varying distances from the astrocytic cell bodies (Fig. 3c-d), which were suggestive of nuclear structures. Topical application of the nuclear dye Hoechst 33342 at 30 dpi confirmed that these structures contained nuclei (Fig. 3c-d and Movie 2), revealing a remarkable proliferative capacity of reactive perilesional astrocytes.

**Fig. 3.**
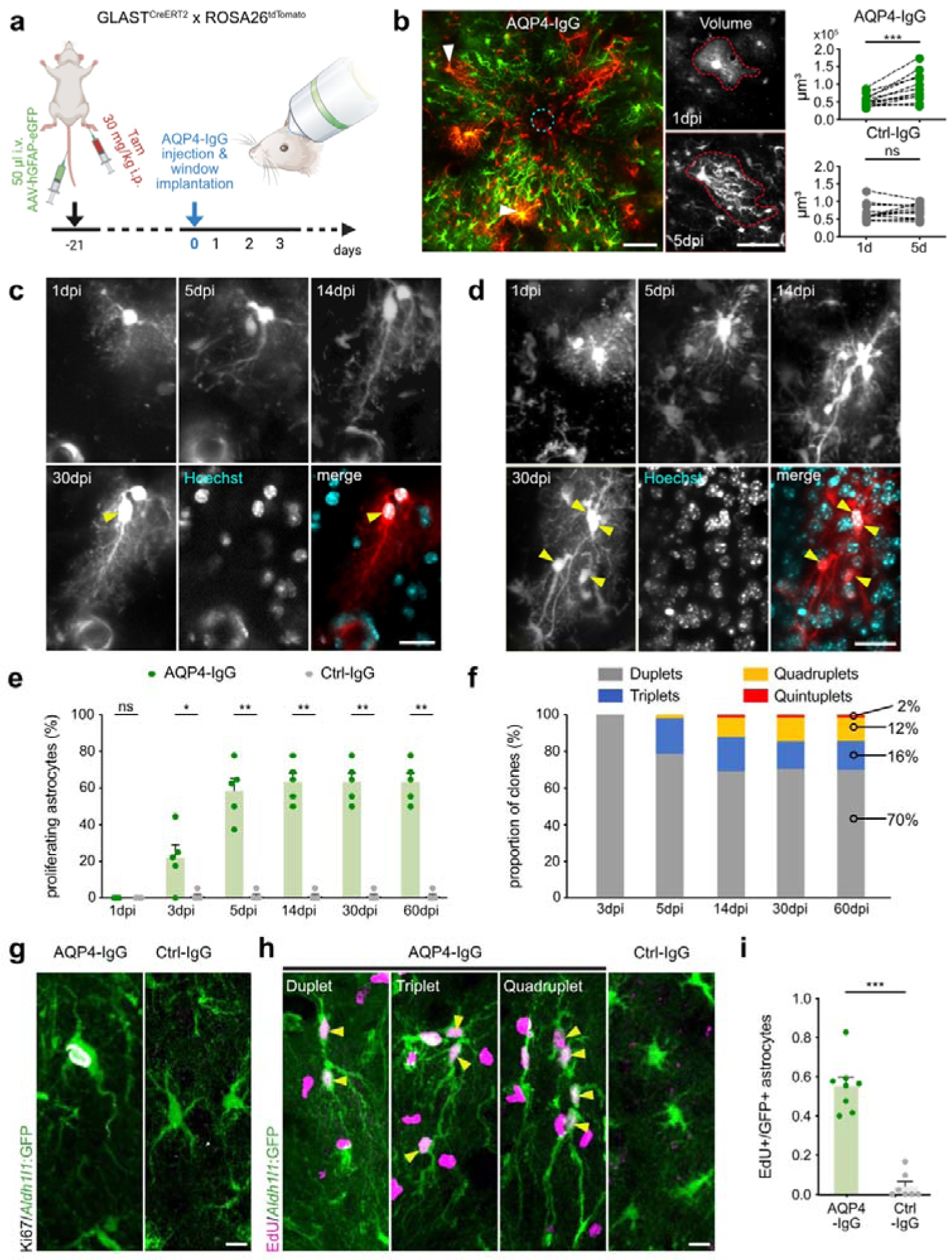
Perilesional astrocytes undergo multiple asymmetric cell divisions. **a**, Sparse-cell labeling protocol for intravital two-photon imaging of individual astrocytes in Glast^CreERT2^ x ROSA26^tdTomato^ mice (Tam = Tamoxifen). **b**, Tracking single tdTom+ astrocytes over time (white arrowheads, left panel, scale bar = 50 µm, cyan dashed circle: injection site) allows detailed morphological evaluation, including astrocytic volume change. Middle panel: z-projected stack of a representative astrocyte volume change (upper cell from left) from day 1 to day 5 (scale bar = 20 µm). Right panels: quantification of volume change in AQP4-IgG versus Ctrl-IgG condition, n = 12 cells from 4 mice, respectively (Wilcoxon signed-rank test). **c-d**, Examples of single perilesional astrocytes and their proliferative activity with one (**c)** or more (**d**) daughter cells (yellow arrowheads) counterstained *in vivo* at the last imaging time point (30 dpi) with the nuclear stain Hoechst 33342 (scale bars = 20 µm; see also Movie 2). **e**, Percentage of proliferating perilesional astrocytes, n = 5 mice for both conditions, AQP4- and Ctrl-IgG (Mann-Whitney test). **f,** Distribution of division cycles in proliferating perilesional astrocytes over time (n = 5 mice). **g-h,** Immunohistochemical staining for Ki67 (**g**) and EdU (**h**) in astrocytes from AQP4- and Ctrl-IgG-injected areas. Yellow arrowheads indicate EdU+ astrocytic clones (scale bars = 10 µm). **i**, Ratio of EdU+ to all GFP-positive perilesional astrocytes in both conditions; n = 7-8 lesions from 6-8 mice per condition (Mann-Whitney test). *p < 0.05, **p < 0.01, ***p < 0.001, ns = not significant. Data represented as mean ± SEM.

With this longitudinal analysis, we could trace all newly appearing intralesional astrocytes back to their corresponding perilesional reactive mother cells and found that approximately 60% of all perilesional astrocytes proliferated within 60 days after the lesion induction (Fig. 3e), compared to only 1% within Ctrl-IgG areas. This proliferation was not restricted to the juxtavascular niche (defined as <5 µm away from a vessel wall). In contrast to previous studies,^4,27^ we observed that mature astrocytes are able to enter the proliferation cycle several times, giving birth to up to 4 daughter cells (Fig. 3f). These results strongly imply asymmetric divisions of perilesional mature astrocytes, which contrasts the prevailing belief that cortical astrocytes proliferate only symmetrically.

Immunohistochemical staining with the proliferation marker Ki67 confirmed astrocytic proliferation (Fig. 3g) in AQP4-IgG compared to Ctrl-IgG areas. However, Ki67 only detects cells that are in active phases of proliferation. To assess the proliferative capacity over a longer period of the repopulation phase, we treated mice with the nucleoside thymidine analog 5-ethynyl-2’-deoxyuridine (EdU) for 7 days after lesion induction. The EdU detection in reactive astrocytes with co-immunolabeling of endogenous Aldh1l1^GFP^ and GFAP in fixed tissue revealed EdU-positive (EdU+) perilesional astrocytes in AQP4-IgG areas (Fig. 3h-i). Again, a variable number of daughter cells could be observed in reactive perilesional astrocytes. In line with the *in vivo* data, around 50-60% of perilesional astrocytes were EdU+ at 14 dpi (Fig. 3i). These results imply a previously unknown proliferative competence of cortical astrocytes in adult brain, which is both independent of the vascular niche and is not restricted to a single cell division.

### Multinucleated astrocytes adopt NPC migratory mechanisms for repopulation

We next wanted to visualize astrocyte invasion dynamics with high temporal resolution *in vivo*. In the adult, cortical astrocytes are considered stationary without the capability to migrate. Upon brain injury, reactive astrocytes stay in their position at the rim of lesion borders, they proliferate and form the glial scar.^4,5,28^

Repetitive imaging of individual proliferating astrocytes revealed daughter cells at large distances away from their mother cells (Fig. 4a-c). The radius of cortical astrocytic domains ranges between 20-30 µm.^29,30^ Based on these previously defined astrocytic domains, we grouped daughter cells into those with a short-range (< 30 µm) and a long-range displacement (> 30 µm). Migration activity was highest during the first 14 days after AQP4-IgG injection (Fig. 4c). Approximately 75% of daughter cells re-localized within the first 7 days, of which >60% (± 4.7) moved more than 30 µm away from their mother cell (Fig. 4d). In all Ctrl-IgG areas only one single cell re-localized.

**Fig. 4.**
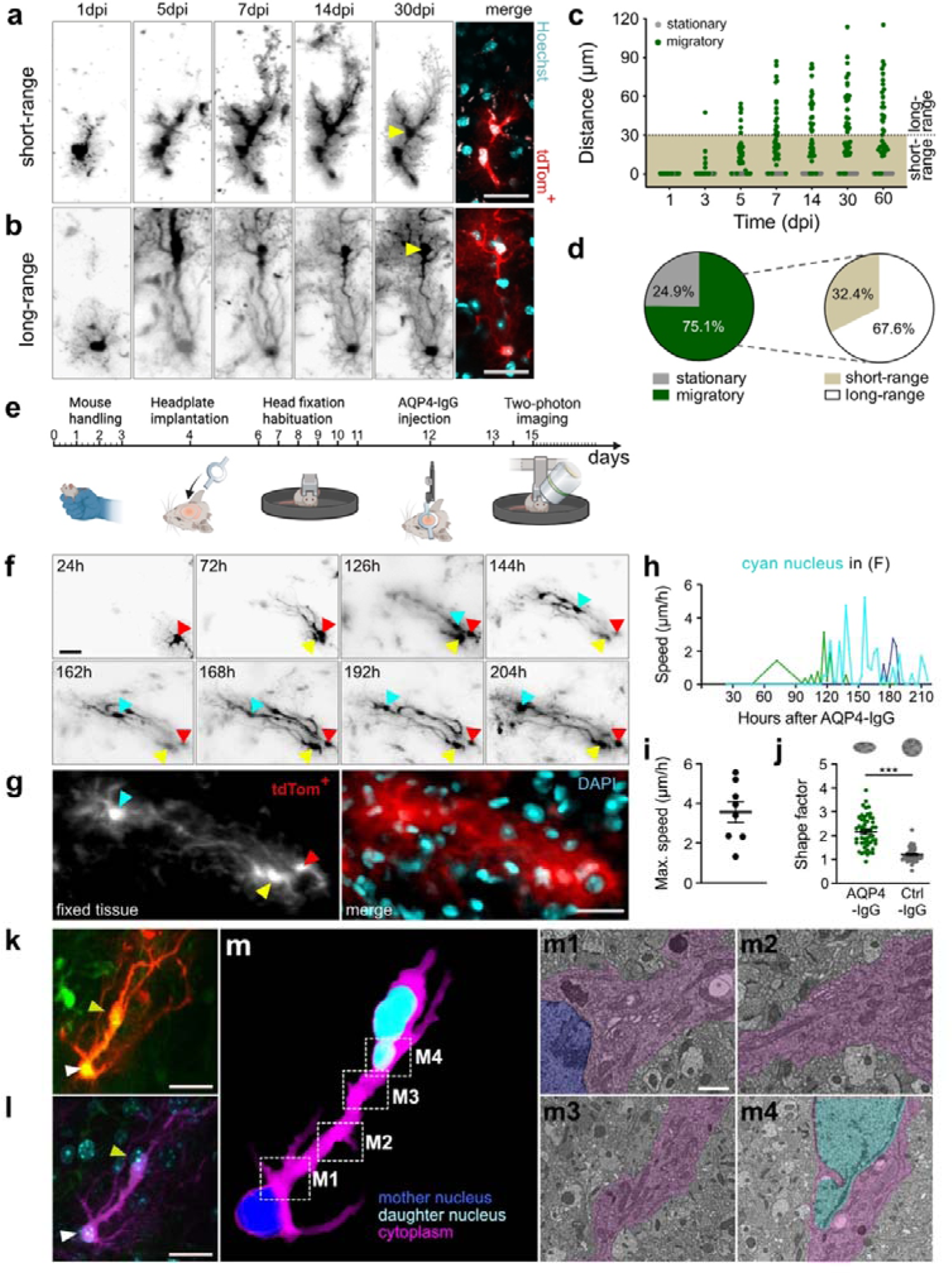
Multinucleated astrocytes use nucleokinesis for lesion repopulation. **a-b**, Examples of tdTom+ astrocytes undergoing division followed by short-range (A) or long-range (B) translocation of daughter cells (yellow arrowheads, scale bar = 20 µm); nuclear counterstaining at last imaging time point (30 dpi) with Hoechst 33342 *in vivo*. **c**, Distance of adult-born astrocytes from their mother cells over time (n = 61 cells from 6 mice). Dashed line indicates the threshold for short- and long-range migration. **d,** Frequencies of different migratory behaviors of reactive proliferating astrocytes (data from c). **e-j,** High-frequency time lapse imaging in awake mice revealing saltatory movement of adult-born astrocytic nuclei. **e**, Experimental workflow; **f**, Time lapses of an individual perilesional astrocyte (red arrowhead; see also Movie 3), showing its reactivation and proliferation, giving birth to two progenies (cyan and yellow arrowheads). Yellow arrowhead indicates the stationary daughter cell and cyan arrowhead shows the migratory daughter nucleus (scale bar = 20 µm). **g**, The astrocyte from **f** and its daughter cell is identified in post-fixed tissue and counterstained with DAPI (scale bar = 20 µm). **h**, Examples of speed traces of individually tracked astrocytic nuclei as well as **i** distribution of speed maxima (n = 7 nuclei from 4 mice). Note the elongated shape of the migratory nucleus during nucleokinesis in **f**. **j**, Change in nuclear shape in astrocytes 5 dpi in AQP4-IgG-(n= 52 nuclei from 5 mice) and Ctrl-IgG injected areas (n = 48 from 4 mice) in fixed tissue, ***p < 0.001 (Mann-Whitney test). Data represent mean ± SEM. **k**, *In vivo* two-photon z-projected stack of a repopulating astrocyte (white arrowhead) with migratory daughter nucleus (yellow arrowhead) 10 days after AQP4-IgG injection (scale bars = 20 µm). **l**, Multi-photon image of the cell pair from (K) in post-fixed tissue, counterstained with DAPI (cyan; scale bars = 20 µm). **m**, 3D surface rendering of the cell pair from **k** and **l** from correlative volume electron microscopy (EM) analysis by tape-based scanning EM. **m1-m4**, Representative EM sections (boxed areas in **m**, demonstrating that mother and daughter nuclei share the same cytoplasm (scale bar = 1 µm). The EM segmentation is representative of at least 3 biological replicates.

Daily two-photon imaging sessions were insufficient to precisely follow the movement pattern and speed of migratory daughter cells. To overcome this limitation, we applied a high-frequency two-photon imaging protocol in awake mice (Fig. 4e). We imaged the mice every 3 hours over 6 consecutive days and precisely captured the dynamics of migratory daughter nuclei (Fig. 4f-h). Strikingly, we observed adult-born daughter cell nuclei being squeezed through process extensions of mother cells in a saltatory manner (Fig. 4f-h and Movie 3). The maximal speed of this process reached 3.5 µm/h with maximal peak rates of 5.6 µm/h and complete stops in between for up to 9 hours. The moving nuclei adopted an ellipsoid shape, regaining their circular form on completion of translocation (Fig. 4f). Analysis of the nuclear shape by DAPI staining in fixed tissue at the time of maximal migratory activity (5-8 dpi) confirmed that the nuclei of intralesional astrocytes had a higher shape factor (length:width ratio) compared to control regions (Fig. 4j). This result is consistent with previously shown morphological changes of nuclei in migratory neural cells during nucleokinesis.^31^

During development, nucleokinesis is a critical part of newborn NPC migration, in which an orchestrated extension of leading processes and movement of cell bodies and nuclei enable guided migration to populate the neocortex.^31^ After separation from the mother cell, NPCs use long radial glial fibers as guidance to find their final location.^32^ We thus wondered whether the observed nucleokinesis in perilesional astrocytes hijacks mechanisms of progenitor-cell migration. Two migration scenarios are conceivable. Either mother and daughter cells separate their cytoplasm, shortly after mitosis, completing the mitotic cell division by cytokinesis with daughter cells migrating afterwards to a new position and including nucleokinesis in a manner similar to newborn neurons. Alternatively, activated astrocytes remain, after mitosis, in a multinucleated intermediate phase, squeezing the daughter cell nuclei to their final destination, and only complete cell division by cytokinesis after nuclear translocation. Given the resolution limits of light microscopy, we used correlative light and electron microscopy (CLEM) followed by serial section electron microscopy to evaluate the ultrastructure of repopulating astrocytes. EM segmentation of dividing perilesional astrocytes with long-range daughter cells and careful tracking of their membranes at 10 dpi revealed that, despite their distance from each other, mother and daughter nuclei share the same cytoplasm (Fig. 4k-m). Daughter cell nuclei again had an elongated shape (n = 3 mice) and sometimes nuclear blebbing (Fig. 4m4), as is known to transiently occur during cell migration of different cell types.^33,34^ Overall, these findings suggest that repopulating astrocytes delay cytokinesis and remain in a multinucleated state for several days. Thus, efficient regeneration after local astrocyte loss appears to rely on the translocation of adult-born nuclei that is guided within the cytoplasmic machinery of the mother cell.

Cell migration requires transcriptional changes and cytoskeletal remodeling for orchestrated translocation. While NPCs are fairly motile cells, they lose their ability to move after differentiation into mature astrocytes.^35^ In our model, we pondered whether perilesional astrocytes reactivate progenitor-like programs, which may explain their migratory behavior. Indeed, immunohistochemical staining for progenitor cell hallmark proteins^11^ revealed a strong upregulation of GFAP (Fig. 2f), Vimentin and Nestin in repopulating astrocytes (Extended Data Fig. 3a-b). Another NPC marker, SOX2, was not upregulated compared to the control at any tested time points (1, 5 and 14 dpi). These results suggest that a subset of reactive astrocytes can indeed regain progenitor-like migratory properties for regeneration after focal astrocyte loss, hereafter referred to as reactive progenitor-like (REPL) astrocytes.

**Extended Data Fig. 3.**
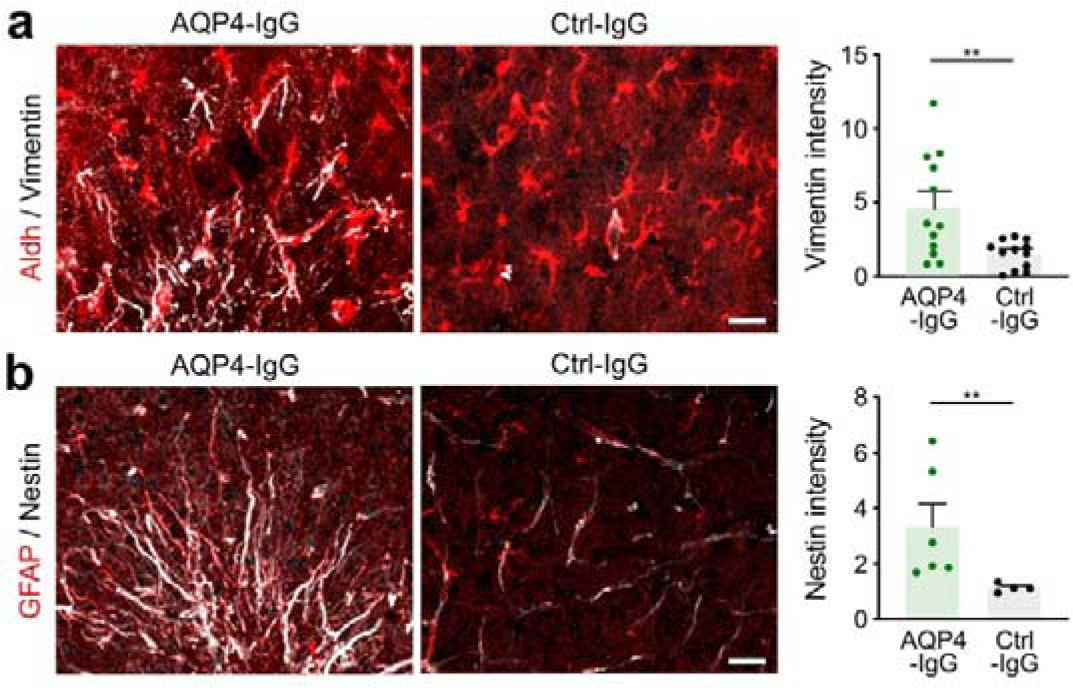
Repopulating astrocytes express progenitor-like hallmark proteins. **a-b**, Left and middle panels: representative confocal images of fixed AQP4-/Ctrl-IgG-injected areas from Aldh1l1^GFP^ mice 14 dpi, stained with the precursor cell markers Vimentin in **a** and Nestin in **b** (in white, scale bar = 20 µm). Note the Nestin-positive vasculature in **b.** Right panels: corresponding quantification of Vimentin and Nestin area intensity, n = 12 lesions from 11-12 mice per condition for Vimentin and n = 4-6 lesions from 4-5 animals for Nestin. **p < 0.01 (Mann-Whitney test). Data represented as mean ± SEM.

### Astrocyte regeneration does not require connexin signaling

During development, glial progenitor cells are known to express different connexins, especially connexin 43 (Cx43),^36^ not only for gap-junctional communication but also for supporting neuronal migration by providing connexin adhesive contacts^37^. In adult brain, mature astrocytes express Cx30 and Cx43 subforms of gap junctions allowing a direct exchange of neurotransmitters and metabolites for intercellular communication.^38^ This prompted us to study the contribution of connexin signaling to REPL astrocyte behavior during regeneration.

We discovered that immunohistochemical analysis of Cx43 expression in perilesional astrocytes in AQP4-IgG areas compared to astrocytes in Ctrl-IgG areas from Aldh1l1^GFP^ mice showed a decrease in the density of Cx43 expression (Extended Data Fig. 4a-b). To further determine a possible role for connexin signaling in REPL astrocytes during lesion repopulation, we used recently generated inducible astrocyte-specific double Cx30 and Cx43 conditional knockout mice,^39^ in which connexins are completely depleted from astrocytes (Extended Data Fig. 4c). Connexin knockout mice (cKO) revealed no difference in lesion size, recovery (Extended Data Fig. 4d), and no difference in percentage of process elongation or polarization (Extended Data Fig. 4e-f). Plus, the number of activated astrocytes and the proliferation rate remained comparable to littermate controls (Extended Data Fig. 4g-h). This makes it unlikely that connexins play a central role in astrocytic regeneration.

**Extended Data Fig. 4.**
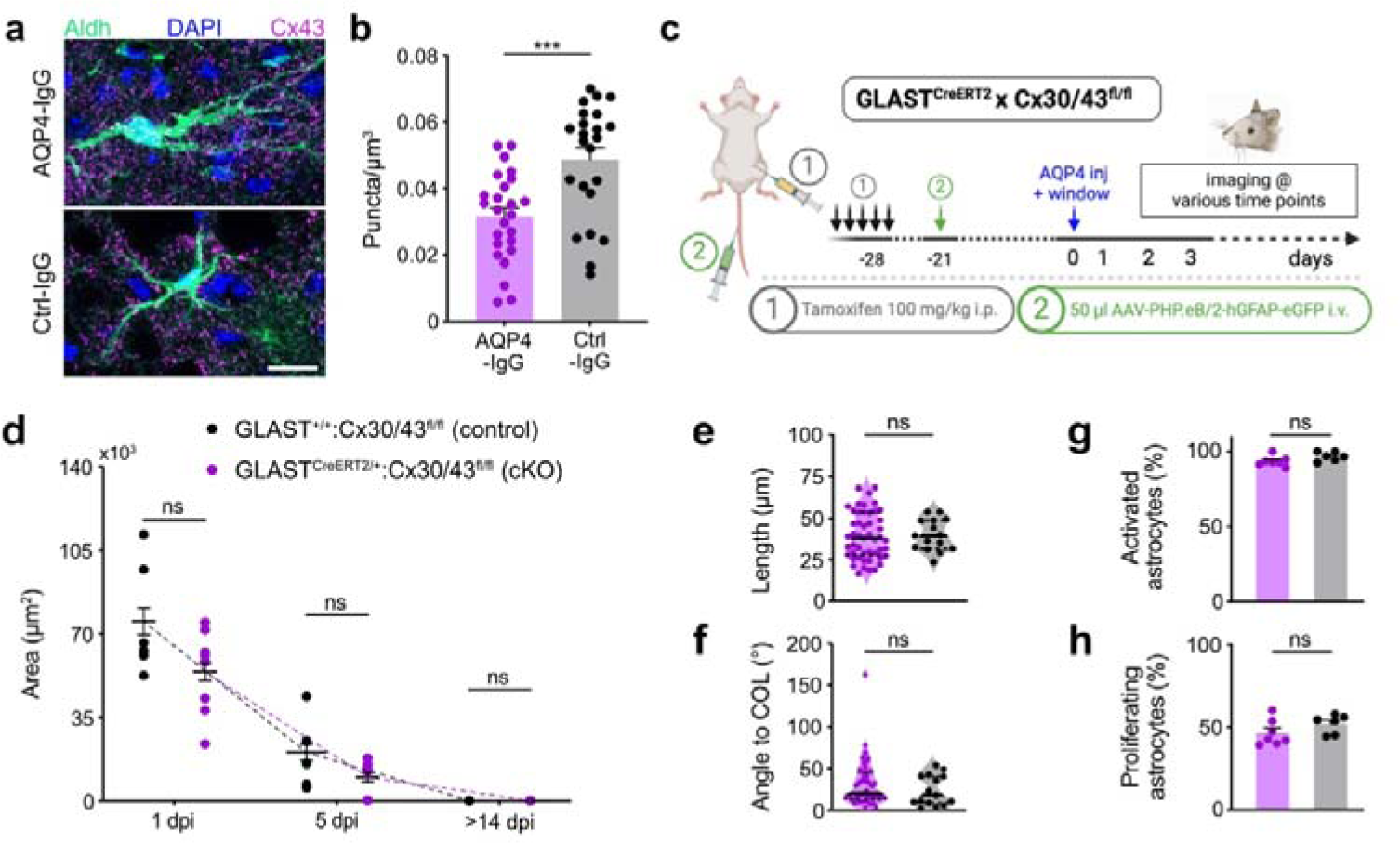
Repopulation of astrocyte-depleted areas is independent of gap junction coupling. **a**, Confocal images of representative astrocytes in AQP4-/Ctrl-IgG-injected areas from post-fixed Aldh1l1^GFP^ mice, stained with Cx43 (scale bar = 10 µm). **b,** Ratio of Cx43 puncta per astrocyte domain from fixed tissue. n = 23-25 cells from 6 animals per condition (Mann-Whitney test). Data represent mean ± SEM. **c,** Experimental approach to study the contribution of astrocyte-specific Cx30 and Cx43 on astrocytic regeneration. **d**, Time course of lesion size from connexin knockout mice (cKO) versus littermate controls *in vivo* after AQP4-IgG injection. n = 8 lesions from 6 cKO mice; n = 6 lesions from 4 control mice. ns = not significant (Mann-Whitney test). Data represent mean ± SEM. **e-f**, Vector-based quantification of process length and polarization of perilesional astrocytes in cKO versus littermate controls. n = 5-12 astrocytes per lesion from 3-7 mice per condition (Mann-Whitney test). Thick black line represents the median and fine black dashed lines represent the first and third quartiles. **g-h**, Percentage of reactive and proliferating astrocytes is comparable between cKO and control mice. n = 6 lesions from 6 cKO vs 6 lesions from 4 littermate control mice (Mann-Whitney test). Data represent mean ± SEM. ***p < 0.001, ns = not significant.

### Astrocyte repopulation is independent of NLRP3

AQP4-IgG-based astrocyte ablation relies on immune-mediated responses. The nucleotide oligomerization domain (NOD)-like receptor family has emerged as a key contributor to the inflammatory environment in different organs.^40^ In the CNS, the NLRP3 subtype has been shown to drive neuroinflammatory responses in animal models of different neurological disorders.^41,42^ We wondered whether the transformation to REPL astrocytes might be triggered by the activation of NLRP3 signaling.

In our model, a global NLRP3 knockout (NLRP3^−/−^) crossed with the Aldh1l1^GFP^ mouse line led to enlarged astrocyte-depleted areas at 1 dpi compared to littermate controls Aldh^GFP^ x NLRP3^+/−^ (Extended Data Fig. 5a-b). The astrocytic lesion was, however, restored by 5 dpi (Extended Data Fig. 5b). We also observed significantly longer astrocytic processes in NLRP3^−/−^ knockouts (Extended Data Fig. 5c). Perilesional astrocytes displayed the same polarization (Extended Data Fig. 5d) but the percentage of proliferating astrocytes significantly increased Extended Data Fig. 5a and 5e). We assume that the increased proliferation compensates for the larger lesion size after astrocyte depletion, rather than being a direct effect of the lack of NLRP3. To test this, we compared the ratio of proliferating astrocytes with the corresponding initial lesion area. Indeed, this analysis revealed that the lesion size correlated with the proliferation activity of perilesional astrocytes (r = 0.90, n = 10 mice, p < 0.001, nonparametric Spearman’s correlation). These findings imply that the NLRP3 pathway does not directly interfere with astrocyte regeneration process, but rather limits the initial AQP4-IgG-mediated astrocyte loss, probably by modulating neuroinflammatory signals of innate immune cells (e.g. microglia, macrophages) that are known to express the NLPR3 inflammasome.^43^

**Extended Data Fig. 5.**
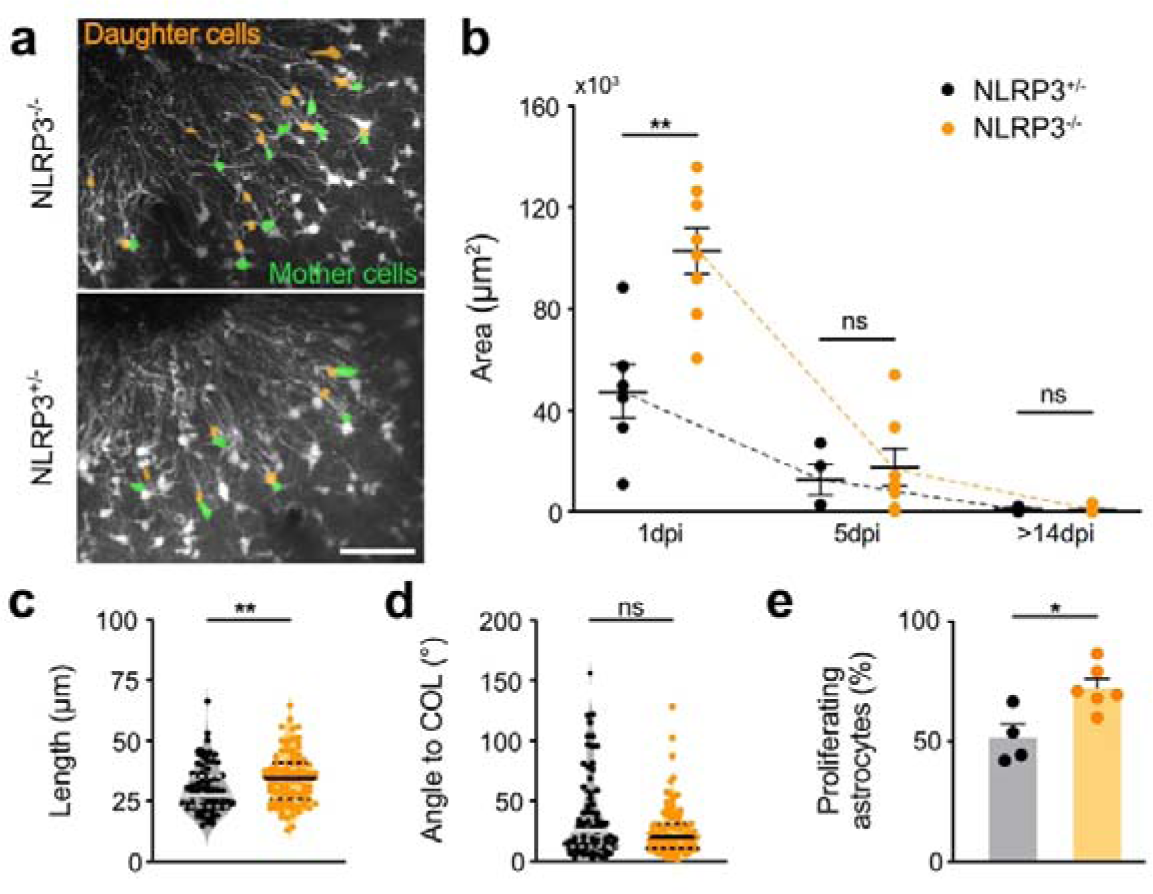
Astrocyte activation upon focal depletion does not depend on NLRP3 signaling. **a**, Two-photon images of perilesional area in Aldh1l1^GFP^ x NLRP3^−/−^ mice compared to heterozygous littermate controls. Note the increased number of proliferating perilesional astrocytes (in green) and their corresponding daughters (in orange) in the knockout animals (scale bar = 50 µm). **b**, Time course of the lesion area size for both genotypes, n = 7-8 lesions from 6 NLRP3^−/−^ mice, n = 4-6 lesions from 4 NLRP3^+/−^ mice (Mann-Whitney test). Data represent mean ± SEM. **c-d,** Vector-based quantification of process length and polarization, n = 92 astrocytes from 4 NLRP3^−/−^ mice, n = 72 astrocytes from 3 NLRP3^+/−^ mice (Mann-Whitney test). Thick black lines represent the median and fine black dashed lines represent the first and third quartiles. **e**, The percentage of proliferative perilesional astrocytes, showing an increase in NLRP3 knockout mice compared to littermate controls, 3-4 mice per condition (Mann-Whitney test). Data represent mean ± SEM. *p < 0.05, **p < 0.01, ns = not significant.

### Ablation volume drives the REPL phenotype

To test whether lesion volume influences the transformation to REPL astrocytes, we chose an approach that fulfills two requirements: i) it causes selective astrocyte ablation in an area of similar size in the same brain region, and ii) it is not associated with the induction of an inflammatory or hypoxic environment resulting from necrotic cell death.

The recently developed 2Phatal method successfully ablates cells through programmed cell death.^18^ This method was originally designed for single neuron/astrocyte ablation. Therefore, we adapted it to encompass large cell numbers (Fig. 5a). In line with previous studies, targeting single astrocytes with 2Phatal induced slow nuclear pyknosis and the formation of apoptotic bodies leading to cell death within 5-10 days.^18^ 2Phatal ablation of a group of 3-5 astrocytes did not show a significant reaction in surrounding astrocytes (Fig. 5b-e) compared to control areas (Hoechst staining, but no laser ablation). Next, we performed ablation of astrocytes covering an area comparable to the mean area of our AQP4-IgG-mediated astrocyte depletion (40000 µm^2^). This led to a prominent reactivity of perilesional astrocytes with process elongation and polarization in only one astrocytic layer (∼30-40 µm in z-dimension) but did not result in substantial proliferative or migratory astrocyte activity (Fig. 5b-e). However, when 2Phatal was performed on several astrocytic domains in all directions (over 100 µm in z-dimension), surrounding astrocytes not only became reactive but also underwent multiple proliferation and developed migratory behavior (Fig. 5b-e). These results indicate that transformation of mature astrocytes into REPL astrocytes does not require a specific neuroinflammatory milieu, but rather represents a conserved fundamental mechanism to restore astrocytic networks when hypertrophy and process elongation of surviving astrocytes alone are not sufficient to compensate for astrocyte loss.

**Figure 5.**
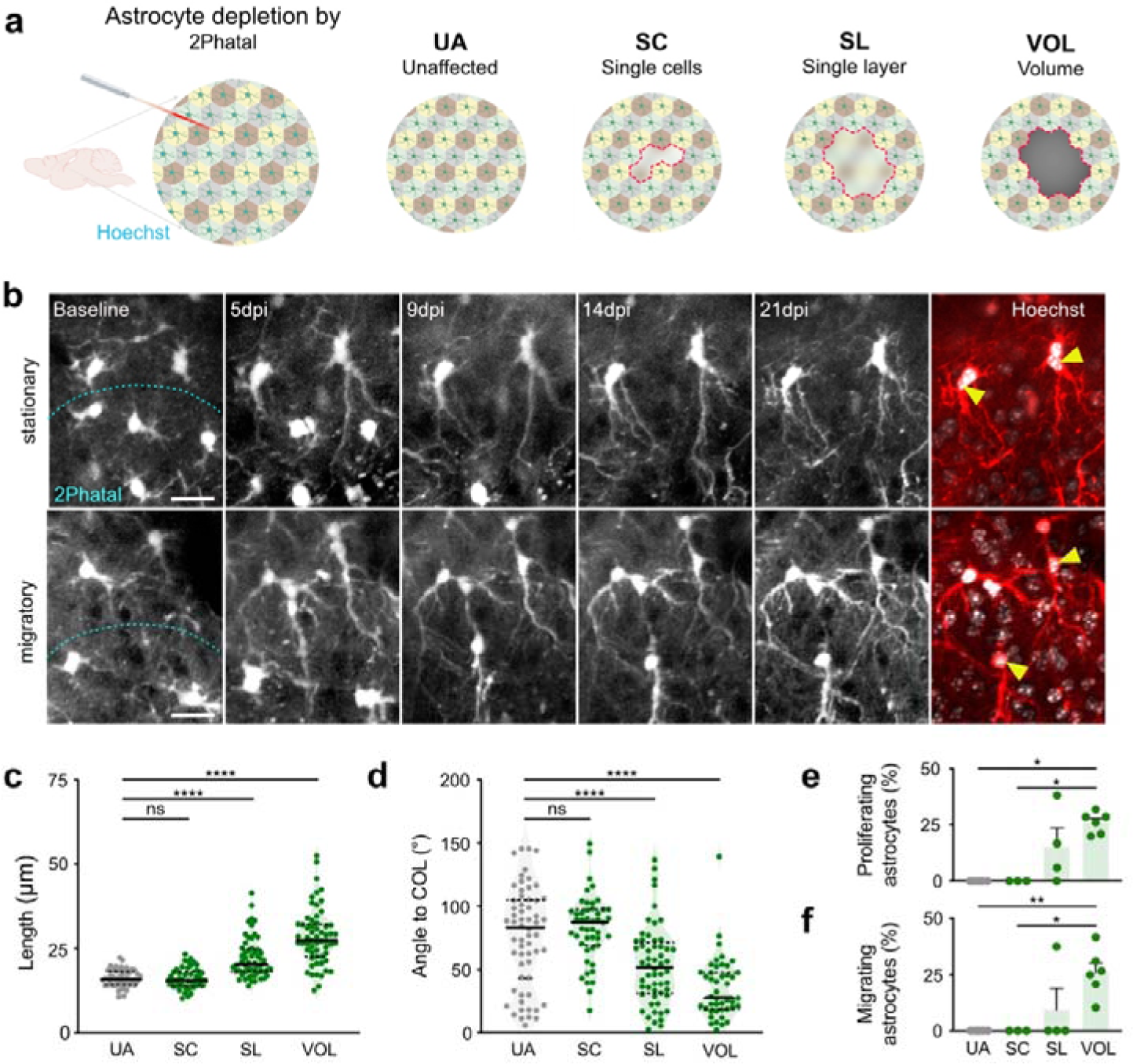
2Phatal targeted astrocyte ablation volume drives the REPL phenotype. **a**, Experimental setup for astrocyte ablation by modified 2Phatal method in different configurations: UA = unaffected area, i.e. area without ablation but with *in vivo* Hoechst staining, SC = single cells (∼5 cells, 0.08-0.14 x 10^6^ µm^3^), SL = single layer (∼1.3-2.2 x 10^6^ µm^3^), VOL = volume ablation (∼6.3-10 x 10^6^ µm^3^). **b**, Representative two-photon *in vivo* time lapse images of perilesional astrocytes 1-21 days after 2Phatal VOL ablation in Aldh1l1^GFP^ mice. Dashed line indicates the border of the 2Phatal ablated area. Note the slow disappearance of 2Phatal targeted astrocytes within two weeks. At the last imaging time point, Hoechst 33342 (in white) was added to counterstain for nuclei (astrocytes are shown in red). Stationary (upper panel) and migratory (lower panel) daughter cell nuclei are indicated by yellow arrowheads. Scale bars = 20 µm. **c-d**, Vector-based quantification of process length and polarization, n = 5-18 astrocytes per mouse, 3-6 mice per condition (Mann-Whitney test). Thick black lines represent the median and fine black dashed lines represent the first and third quartiles. **e-f**, The percentage of proliferative and migrating perilesional astrocytes, respectively, showing an increase with larger ablation volumes, 3-6 mice per condition. Data represent mean ± SEM. *p < 0.05, **p < 0.01, ****p < 0.0001, ns = not significant.

### REPL phenotype is present in human astrocytopathy

Having identified REPL astrocytes after AQP4-IgG-mediated astrocyte demise in the murine brain, we asked whether similar astrocytic phenotypes are present in human AQP4-IgG-mediated lesions. For this purpose, we examined human brain tissue from confirmed cases of AQP4-IgG positive NMOSD (Extended Data Table 1). Lesions were identified by AQP4 protein loss (Fig. 6a) and by marked GFAP upregulation (Fig. 6b) at lesion borders. Indeed, a subset of reactive astrocytes in the perilesional region showed a multinucleated state with elongated and polarized shapes (Fig. 6d-g and 6i) that were significantly more abundant than in non-affected areas (Fig. 6c and 6i). Moreover, some perilesional astrocytes were positive for the proliferation marker Ki67 (Fig. 6h). These results suggest that REPL astrocytes are also present in human AQP4-IgG-mediated pathologies and potentially contribute to their lesion recovery.

**Fig. 6.**
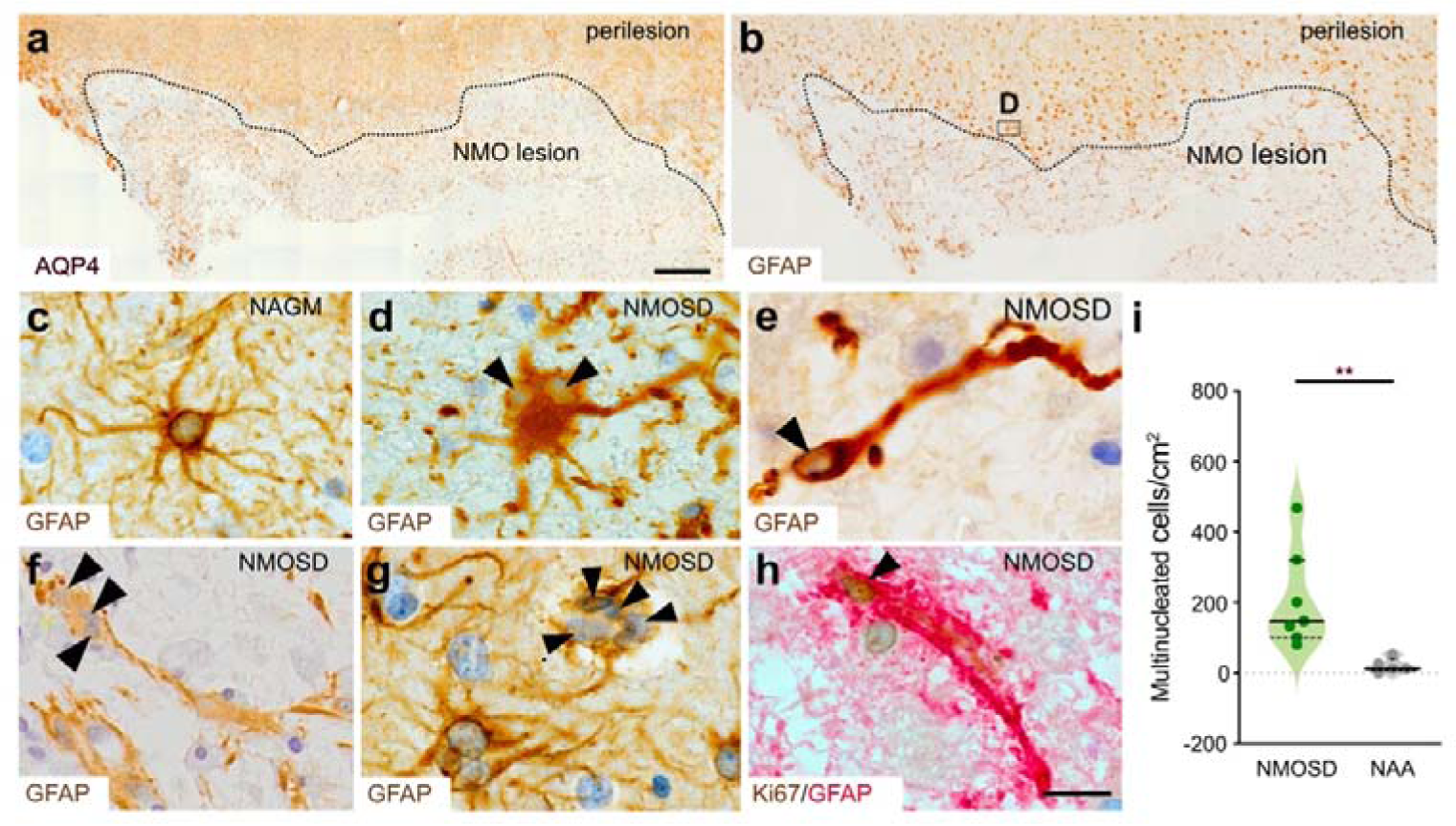
REPL phenotype in human AQP4-IgG-mediated NMOSD lesions. **a-b**, Human brain biopsy, showing a NMOSD lesion and perilesional area stained against AQP4 in **a** and GFAP in **b** with marked loss of AQP4 and GFAP within the lesion. Dashed line indicates the border of the lesion (scale bar = 200 µm). Boxed area magnified in **d**. **c**, GFAP staining of normal-appearing gray matter (NAGM), illustrating a normal cortical astrocyte. **d-g**, Examples of morphological features of reactive perilesional NMOSD astrocytes stained for GFAP, demonstrating: **d**, diploid astrocyte, **e**, polarized shape, **f**, triploid cell with a leading polarized process, **g,** tetraploid cell. Black arrowheads indicate individual nuclei. Scale bar = 10 µm. **h**, Example of a reactive polarized perilesional astrocyte stained for GFAP (in red), showing Ki67 positivity (in brown, indicated by arrowhead). Scale bar = 10 µm. **i**, Quantification of the number of multinucleated astrocytes per cm^2^ found in NMOSD perilesional areas (n = 7) compared to non-affected areas (NAA; n = 5). Nuclear counterstaining with hematoxylin in panels **a-h**. Thick black line represents the median and fine black dashed lines represent the first and third quartiles. **p < 0.01 (Mann-Whitney test).

**Extended Data Table 1.**
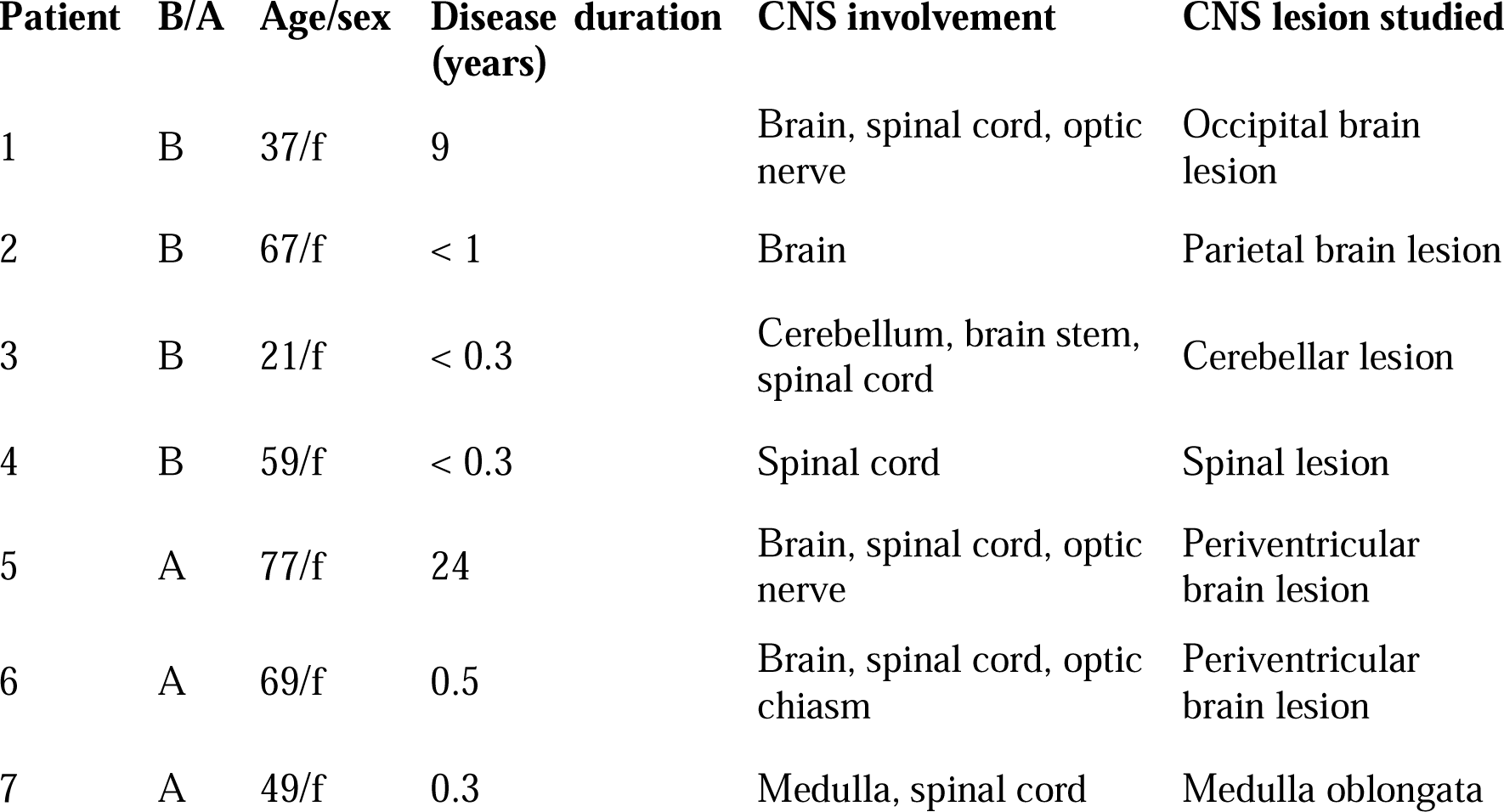
Clinical data of patients included in study.

| Patient | B/A | Age/sex | Disease duration<br>(years) | CNS involvement | CNS lesion studied |
| --- | --- | --- | --- | --- | --- |
| 1 | B | 37/f | 9 | Brain, spinal cord, optic nerve | Occipital brain lesion |
| 2 | B | 67/f | < 1 | Brain | Parietal brain lesion |
| 3 | B | 21/f | < 0.3 | Cerebellum, brain stem, spinal cord | Cerebellar lesion |
| 4 | B | 59/f | < 0.3 | Spinal cord | Spinal lesion |
| 5 | A | 77/f | 24 | Brain, spinal cord, optic nerve | Periventricular brain lesion |
| 6 | A | 69/f | 0.5 | Brain, spinal cord, optic chiasm | Periventricular brain lesion |
| 7 | A | 49/f | 0.3 | Medulla, spinal cord | Medulla oblongata |

## DISCUSSION

We here developed two different methods of selective astrocyte ablation, accessible to chronic *in vivo* two-photon microscopy. We discovered that in response to focal astrocyte loss, surviving perilesional cortical astrocytes develop a remarkable, so far unknown, plasticity for efficient lesion repopulation. To achieve this, mature astrocytes transform into reactive progenitor-like (REPL) cells, exhibiting drastic morphological changes, transient calcium hyperactivity, asymmetric cell division, a multinucleated interstage, and migratory behavior of their daughter cells via nucleokinesis. We show that the REPL phenotype does not require a neuroinflammatory trigger but is mainly driven by the volume of astrocyte loss.

The ability of astrocytes to proliferate has previously been described following brain and spinal cord injury.^4,5,11,12^ However, our results question the prevailing assumptions that astrocyte proliferative programs require disruption of the BBB, are restricted to the juxtavascular niche, are limited to one cycle of symmetric cell division and are characterized by their non-migratory nature. On the contrary, we were impressed to see that vessels with a complete loss of astrocyte endfeet coverage do not exhibit any immediate alteration in their volume fraction, blood flow or diameter under basal conditions *in vivo*. The functional analysis of single capillaries in AQP4-IgG-mediated lesions suggests that the gross vascular integrity remains intact. We also found no indication of a BBB leakage 1 day after lesion induction, measured by cadaverine extravasation (data not shown). However, subtle deficits in blood-brain-barrier integrity cannot be completely excluded.^44^

In the developing brain, neural- and glial-lineage progenitors migrate into the cortex along radial glial fibers.^35,45^ Notably, previous *in vivo* studies of CNS pathology did not find direct evidence of cortical astrocyte migration in the adult brain in response to CNS insults.^4–6^ Using intravital imaging with high temporal resolution, we here provide compelling evidence that a subset of reactive astrocytes can repopulate focal lesions by activating NPC-like mechanisms leading to multiple cell divisions and migration. One possible reason why REPL cells have so far not been discovered could be due to the fact that more unspecific lesions cause the formation of a glial scar, which might impede the access of resident cells into the lesion core.

Within the CNS, nucleokinesis has so far mainly been described in NPCs during development.^31^ These studies unraveled the involved dynamics of cytoplasmic organelles and cytoskeletal components. Indeed, the REPL astrocyte nucleokinesis shows features that have been described in the migration process of NPCs in mouse embryo.^46^ Being an ATP-consuming saltatory movement with rapid extension of the leading process and forward displacement of the nucleus, nucleokinesis is a challenge for the large size of the nucleus and requires a remodeling of its shape and stiffness,^47^ which explains the transient shape change of the travelling nuclei seen in our model.

In previous *ex vivo* studies the migration speed of neural precursors ranged from 20-60 µm/h^32,48^ and depends on neuronal subtype, migration phase and used techniques. *In vivo* imaging data of neural progenitor migration are sparse. Neural precursor migration velocities in anesthetized embryos are much slower, averaging 10-13 µm/h.^35,46^ Data on dynamics of progenitor cell migration in awake adult mice are missing and our *in vivo* two-photon imaging approach closes this gap as it allows a direct observation of migration dynamics. Notably, the speed of REPL astrocytes was considerably slower than cellular migration during development.

Astrocyte reactivity has so far mainly been investigated as a secondary response to injury (e.g. trauma, stroke, toxic substances), to neuroinflammatory conditions or in the context of genetic astrocytopathies (e.g. Alexander disease).^1^ According to this literature, astrocyte regeneration in the adult brain requires a strong neuroinflammatory stimulus and involves connexin signaling. Surprisingly, we found that REPL astrocytes neither required the NLRP3 inflammasome, an important mediator of neuroinflammation in the brain, nor astrocyte-specific connexin signaling. Instead, based on our optical astrocyte depletion protocol we conclude that the volumetric extent of astrocyte loss is the key driver of the REPL astrocyte phenotype. Although the rate of cell death in 2-Phatal induced ablations is slower than the antibody-mediated depletion (days vs. hours), we were able to evoke the same proliferative and migratory behavior. Process length and proliferation rate of REPL astrocytes strongly correlated with lesion volume. An increasing volume of astrocyte depletion most likely exceeds the capacity of the perilesional astrocytes to close the void by hypertrophy and process elongation alone. This discovery challenges our current understanding of reactive astrocytes. Astrocytes seem to have a fundamental ability to regenerate by reactivating NPC-like programs that are regularly reeled off during development and likely involve contact-inhibition pathways for such a syncytial cell type. As this form of astrocyte plasticity is independent of the underlying cause, it is likely to be present in brain pathologies that involve focal astrocyte loss. However, the exact mechanisms remain unclear and further work is required.

During development, migrating NPCs separate shortly after mitosis and migrate further as complete cells. We did not observe a separation between the mother and daughter cell during nucleokinesis, but instead identified a multinucleated interstage, which at least can last for more than one week. While this phenomenon has not been reported before in rodent brain, multinucleated astrocytes are regularly observed in human neurological diseases.^49^ Such regenerative mechanisms in human NMOSD pathology have not been reported, although bipolar GFAP-positive cells have been found in NMOSD lesions and considered as astrocyte precursor cells of unknown origin.^50^ Accordingly, our histopathological analysis showed multinucleated and elongated peri-und intralesional astrocytes in human NMOSD lesions. This favors the existence of REPL astrocytes in the adult human brain.

In conclusion, using chronic *in vivo* imaging we have discovered a hitherto unknown cellular plasticity of mature astrocytes that underlies the effective repopulation of focal glial lesions in the adult brain. Perilesional reactive progenitor-like (REPL) astrocytes can be defined by asymmetric divisions, a prolonged multinucleated interstage and migration via nucleokinesis.

## Methods

### Animals

Male and female 4-to 8-month old mice were used in this study. All animals were kept in temperature- and humidity-controlled husbandry conditions (22-24 °C, 50-60% relative humidity) with unlimited access to food and water. Male and female littermates were equally allocated into control and experimental groups if not explicitly mentioned otherwise. All animal experiments were approved by the local veterinary authorities according to the guidelines of the Swiss Animal Protection Law, Veterinary Office, Canton Zurich (Animal Welfare Act of 16 December 2005, and Animal Protection Ordinance of 23 April 2008).

To visualize astrocytes, the aldehyde dehydrogenase 1 family, member L1 Aldh1l1^GFP^ mouse strain was obtained from MMRRC (Tg(Aldh1l1-EGFP)OFC789Gsat/Mmucd). For experiments measuring astrocytic calcium activity, C57BL/6J mice from Charles River were injected with viral constructs to induce GCaMP6s expression in astrocytes (for details see below). To trace individual astrocytes, GLAST^CreERT2^ x ROSA26^tdTomato^ mice were injected once with a low dose of tamoxifen (30 mg/kg i.p., 3mg/ml in corn oil) to obtain sparse red astrocyte labeling. Additionally, ssAAV-PHP.eB/2-hGFAP-EGFP-WPRE-hGHp(A) virus (50 µl of 2.0 x 10^13^ particles/ml, Viral Vector Facility, UZH, Zurich, Switzerland) was i.v. injected to obtain widespread astrocytic GFP expression for identification of the perilesional area.

GLAST^CreERT2^ x Cx30/Cx43^fl/fl^ mice^39^ were used to study the role of astrocytic gap junction coupling 4 weeks after tamoxifen injection (for 5 consecutive days, 100mg/kg i.p., 10mg/ml in corn oil). Littermates negative for Cre served as control mice (GLAST^+/+^ x Cx30/Cx43^fl/fl^). Over 90% loss of Cx30 and Cx43 in the cortex has been previously confirmed in immunohistochemical sections of these knockout mice at 30 dpi.^39^ The ssAAV-PHP.eB/2-hGFAP-EGFP-WPRE-hGHp(A) was i.v. injected (50 µl of 2.0 x 10^13^ particles/ml) 3 weeks before surgery for visualization of astrocytes. In experiments investigating the role of NLRP3, Aldh1l1^GFP^ mice were crossed with NLRP3^−/−^ mice (JAX stock #021302)^51^.

### Surgical interventions

Surgical interventions were performed on two separate days. During anesthesia, Vitamin A eye ointment was applied to prevent corneal desiccation. To prevent dehydration, 10ml/kg of prewarmed Ringerfundin was subcutaneously injected before beginning the intervention. Oxygen was delivered (200 cc/min) to prevent hypoxemia. Animalś vital signs and temperature were monitored and controlled via MARTA Pad (Vigilitech).

#### Head plate implantation

Animals were anesthetized with 2–2.5% isoflurane in a mixture of O_2_ and air (30%/70%) at a flow rate of 300 ml per min. For head plate implantation, animals were fixed in a stereotaxic frame (Model 900; David Kopf Instruments). The head region was shaved, disinfected (Kodan; Schülke & Mayr), and anesthetized by local s.c. injection of a mixture of lidocaine (10mg/ml) and bupivacaine (5mg/ml). The skull surface was exposed by a midline incision (1.2 to 1.5 cm long), cleaned by removing the connective tissue and covered with a bonding agent (One Coat 7 Universal, Coltene). Next, a custom-made stainless steel head plate was positioned centrally above the exposed bone and attached by multiple layers of blue light-curing dental cement (Tetric EvoFlow, Ivoclar Vivodent) while sparing the area designated for later craniotomy.

#### AQP4-IgG/Ctrl-IgG injection

2 to 3 days after head plate implantation, animals were anesthetized by i.p. injection of a mixture of fentanyl (0.05 mg/kg bodyweight (BW); Sintenyl; Sintetica), midazolam (5 mg/kg BW, Roche) and medetomidine (0.5 mg/kg BW, Orion Pharma) and reapplied as needed (after about 60 minutes). A craniotomy above the left somatosensory cortex was performed using a dental drill (diameter 0.2 mm, H-4-002HP, Rotatec GmbH). After craniotomy and durotomy, a mixture of a human IgG1 recombinant antibody AQP4-IgG (clone 7-5-53, 0.1-3 µg/µl), reconstructed from a clonotypic plasma blast obtained from the cerebrospinal fluid of a AQP4-IgG-positive NMOSD patient^52^ together with human complement (15-30 U/ml, Sigma-Aldrich, #S1764) was injected using a custom-made microinjector and a pulled glass capillary (Drummond PCR micropipettes, Drummond Scientific) with a tip diameter of 35-45 µm. A volume of 130 nl was distributed between 0-300 µm depth in 100 µm steps (Fig. 1a) and additional 90 nl were slowly infused (20 nl/min) during insertion and retraction of the pipette to keep it patent. In control lesions, the recombinant Ctrl-IgG (0.1-3 µg/µl, clone ICOS-5-2), of the same isotype as AQP4-IgG (human IgG1), of unknown specificity from a meningitis patient was injected together with complement in the same fashion^52^. For two-photon imaging, a chronic window was implanted (see below). Anesthesia was reversed by i.p. injection of a mixture of flumazenil (0.9mg/kg BW) and atipamezole (45 mg/kg BW). After surgeries, animals were treated with buprenorphine (0.1 mg/kg BW s.c.) and carprofen (10 mg/kg BW s.c.). Thereafter, carprofen treatment was continued twice a day for three days.

#### Injection of genetically encoded calcium indicators

In C57BL/6J mice, headplate implantation, craniotomy and durotomy were performed as described above. Three to four neighboring injections of adeno-associated virus (1.6 × 10^12^ particles/mL of AAV9-hGFAP-GCaMP6s; Viral Vector Facility, UZH, Zurich, Switzerland; 125 nl per injection) were performed with a custom-made microinjector to achieve multiple overlapping spots where astrocyte depletion was later induced (see above). Viral injections were performed via a glass capillary at a depth of 350 and 200 μm from the brain surface. Large blood vessels were avoided to prevent bleeding and the absorption of light by hemoglobin during imaging. A chronic window was implanted (see below). Three weeks after virus injection and baseline imaging, the chronic window was again removed and AQP4-IgG/Ctrl-IgG/complement was injected (as described above).

#### Chronic window implantation

For chronic two-photon measurements a square sapphire glass (3 × 3 mm, HEBO Spezialglas) was implanted after the completed virus or antibody injection. The glass was gently placed and sealed with light-curing dental cement (Tetric Evoflow, Ivoclar Vivodent).

#### *In vivo* nuclear staining

In a subset of animals, the cranial window was removed 30 days after lesion induction and the nuclear dye Hoechst 33342 (0.1 mg/ml, 2×15 min) was applied for nuclear staining *in vivo*. After having implanted a new chronic window, lesions were imaged on the following day.

### Behavior training for awake two-photon imaging

Mouse handling started before head plate implantation, multiple times a day for three days, in order to familiarize the animals with the experimenter. Two days after head plate implantation, handling was resumed. Animals were adapted to the head fixation by restraining them with the implanted head plate several times a day, with a gradual increase in restraint time from seconds up to several minutes. The last few training sessions took place in the microscopic setup with the microscope and shutters operating to familiarize the animals with the setup. After an extensive training period of about 1 week, animals tolerated head fixation for the duration of a 15 to 20 min imaging session while being able to move on a custom-made air-lifted platform.

### Anesthetized and awake *in vivo* two-photon microscopy imaging

Two-photon imaging was performed with a custom-made two-photon laser scanning microscope ^53^ equipped with two two-photon lasers (Chameleon Discovery NX, Coherent) and a 25x water-immersion objective (W Plan-Apochromat 25x/1.05 NA, Olympus). The tunable laser was set to 920 nm to excite GFP and GCaMP6s or to 780nm to excite Hoechst.

A second laser with a fixed 1040 nm wavelength line was used to excite tdTomato. The emitted light was focused on the photomultiplier (H9305-03, Hamamatsu) equipped with emission filters for blue fluorescence (475/64 Brightline HC; AHF Analysentechnik), green (535/50 Brightline HC, AHF Analysentechnik) and red fluorescence (607/70, 607/70 Brightline HC, AHF Analysentechnik). A dichroic mirror at 506 nm and 560 nm separated the emission light beam. ScanImage^54^ and custom-written LabVIEW software were used for image control and data acquisition.

#### Anesthetized imaging

mice were anesthetized using isoflurane. During image acquisition, isoflurane was supplied via a face mask (1.5 −2% in O_2_ and air). 24 hours after antibody injection, the first images were acquired and then afterwards at predefined time points (see Results). Stacks were acquired with 512 x 512 pixels at 0.74 Hz for overview and with 2048 x 2048 pixels at 0.37 Hz for detailed signal analysis. For calcium measurements, we chose a concentration of isoflurane that has previously shown very little interference with baseline calcium activity in healthy astrocytes^55^. Calcium signals were measured at frequencies of 11.84 Hz over a period of 120 s (128 x 128 pixels, 0.5 ms/line, zoom 6) from 2 to 3 different spots of astrocyte-depleted or control lesions.

#### Vasculature imaging

the vasculature was labeled by i.v. injection of 50 μl of 2.5% (w/v in saline) 70 kDa Texas Red dextran (Thermo Fisher; D1830). The dye was excited at a wavelength of 870 nm. For structural assessment of the capillary bed, z-stacks were acquired at 1.0 μm step size with a resolution of 512 x 512 pixels at 0.74 Hz. Red blood cell velocity was measured along the vessel midline with line scans (256 x 256 pixels, 6.1 Hz) and subsequently processed by a custom-designed image processing toolbox (Cellular and Hemodynamics Processing Suite^56^ in MATLAB (R2017b; MathWorks) using the Radon transform. Using a Gaussian fitted intensity profile drawn perpendicular to the vessel midline, the vessel’s diameter was calculated at full-width half maximum.

#### Awake imaging

24 hours after antibody injection, initial images were acquired. After 3 to 5 days, acquisitions were performed every 3 hours for 6 consecutive days, followed by imaging sessions at lower frequency as for anesthetized imaging (2, 4 and 8 weeks).

### 2Phatal astrocyte ablation

Headplate implantation, craniotomy and durotomy was performed in Aldh1l1^GFP^ animals as described above. Hoechst 33342 (0.1 mg/ml, Invitrogen) was bath applied topically for 2 x 5 min. Hoechst was thoroughly washed from the cortical surface with Ringer’s solution and a cranial window was put in place as described above. 1 day after surgery, fluorescently-labeled nuclei of different numbers of astrocytes were targeted over a square region of interest (ROI) with a focused two-photon laser (Chameleon Discovery NX, Coherent) tuned to a wavelength of 775nm. ROIs were scanned 240 times with 16 x 16 pixels at 23.67 Hz (resulting in 10 seconds total irradiation time) through a 25x water-immersion objective (W Plan-Apochromat 25x/1.05 NA, Olympus) at 50 mW laser power (measured under the objective) to induce apoptosis (as described in ^18^). Laser output power was increased by 10 mW for every 30 µm increase in z-direction for cells in deeper areas to account for light scattering and absorption. Images of the lesion area were acquired under isoflurane anesthesia at different time points after 2-Phatal cell ablation.

### Image analysis

Images were processed using the open-source image analysis software FIJI (ImageJ 2.1.0).^57^ For display purposes, two-dimensional (2D) images were generated using z-maximum intensity projections for merged channels. In non-quantitative panels, the gamma value was adjusted non-linearly to enhance visibility of low-intensity objects. Datasets were processed with Excel (Microsoft Corporation). All figures were crafted with Affinity Designer 2 (Serif). Graphical illustrations of experimental protocols were generated with BioRender.

#### Vector analysis

Astrocytes were chosen randomly from perilesional areas. Distal ends of primary astrocytic processes, location of cell soma as well as the center of the lesion were manually marked by single-point ROIs in two-photon image stacks taken 1, 5, 14 and 60 days after AQP4-IgG-mediated lesion induction. Coordinates of ROIs in x, y and z were extracted. From these coordinates, every process was assigned a vector that reflected its length and orientation. This allowed for calculation of an average vector for every cell (see Equation 1) by means of a custom written Python script (Python 3.6.5), summarising both length and orientation of the cell’s primary processes. These measures could then be compared over different timepoints. In controls, the average diameter of AQP4-IgG-mediated lesion 1 dpi was chosen as a safety margin to avoid the analysis of possible astrocytic changes in close proximity to mechanical injury. Only astrocytes outside of but adjacent to this margin were

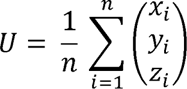

***Equation 1:** Calculation of the coordinates of the average vector U by averaging x, y, and z coordinates of all vectors of a cell.*

#### Lesion size and cell number analysis

*In vivo* image stacks of Aldh1l1^GFP^ mice taken 1 day after antibody injection were z-projected over 50-150 µm depth and used for lesion area calculation. The number of intralesional astrocytes was assessed by manually counting all GFP positive cell bodies that appeared within this lesion area over different imaging timepoints.

#### Proliferation rate

All perilesional astrocytes of a randomly chosen quadrant of the lesion were marked in images taken 1 dpi and then tracked over several time points (up to 60 dpi) to detect proliferation. In control conditions, astrocytes at the same average distance from the site of injection were tracked similarly in random quadrants.

#### Calcium activity

Image analysis was performed using ImageJ and a custom-designed image processing toolbox, Cellular and Hemodynamic Image Processing Suite (CHIPS^56^) based on MATLAB (R2017b, MathWorks). Calcium transients were quantified by an unbiased algorithm^24,58^. ROIs were selected by combining hand-selection of astrocytes based on the based algorithm^58^ using the 128 x 128 pixel images. A 2D spatial Gaussian filter (**σ** = 2 µm) high-resolution acquisition (512 x 512 pixels) and automated ROI detection with an activity- and a temporal moving average filter (width = 1 s) were applied to all images to reduce noise. A moving threshold for each pixel was defined in the filtered stack as the mean intensity plus five times the standard deviation of the same pixel in the preceding three seconds. Using this sliding box-car approach, active pixels were identified as those that exceeded the threshold. Active pixels were grouped in space (radius = 2 µm) and time (width = 1 s).

#### Volume analysis

Glast^CreER^ x ROSA26^tdTomato^ cells in *in vivo* images were segmented using the FIJI Segmentation editor plugin on multiple slices in a z-stack. ROIs were interpolated between slices and the ROI area was determined for every slice. Volumes were compared between images acquired 1 dpi and 5 dpi.

#### Nuclear shape factor calculation

DAPI stainings in fixed sections from AQP4-IgG and Ctrl-IgG injected areas 5 dpi were analyzed. All astrocytes within the lesion were included and the shape factor (length:width ratio) of their nucleus was calculated for each cell.

#### Interventional experiments

In NLRP3 and connexin knockout mice, the analysis of lesion size, vector-based process length, orientation, and proliferation rate between different genotypes was performed in a blinded manner.

### Vessel segmentation analysis

Three-dimensional (3D) volumes were segmented by slicing the volume along the depth direction into 2D slices. To avoid bias in the segmentation algorithm towards extreme values, we set all values larger than the 99^th^ percentile intensity to this cut-off value. To obtain the threshold value for segmentation, we used Li pattern recognition,^59^ implemented in the Python scikit-image library.^60^ To prevent extreme threshold values of the Li algorithm in the presence of imaging artifacts, we set the segmentation threshold to the mean value of the 3D volume if the obtained threshold is less than this for the respective 2D slice. To remove noise and small pixel clusters, one binary opening operation was performed after thresholding. To filter out superficial (0-20 μm depth) large vessels from the 3D volumes, connected areas that covered more than 200,000 voxels were disregarded. The vessel volume was calculated by multiplying the number of voxels classified as vessels with the physical volume of a voxel. The volume fraction was calculated by dividing the number of voxels classified as vessels by the total number of voxels in the ROI. To obtain the vasculature surface area we used the findContours function of OpenCV.^61^ The vessel diameter was calculated by skeletonizing the segmentation^62^ and applying a median filter of voxel size three. After sampling points along the centerline and calculating the distance between each point and the inverse segmentation, the minimum calculated distance was used as the radius of a vessel at a respective point.

### Immunohistochemistry and histological analysis

Animals were i.p. injected with pentobarbital (200 mg/kg BW) for terminal anesthesia. Animals were perfused transcardially with 4% paraformaldehyde (PFA) in 0.0.1 M phosphate-buffered saline (1 × PBS; in mM: 1.5 KH_2_PO_4_, 2.7 KCl, 8.1 Na_2_HPO_4_ and 137 NaCl), postfixed for 3 hours in 4% PFA, then dissected and transferred to 30% sucrose (Sigma) solution in PBS for cryoprotection overnight at 8 °C. Brains were placed in embedding medium (NEG-50®, Richard-Allen Scientific) with the apical cortical surface facing the cutting plane and then cut into 30 μm sections in a cryostat (LEICA CM3050S, Leica Biosystems AG). Antibodies were diluted in 0.3% Triton X-100 (Sigma) and 5% normal donkey serum (ab7475, Abcam) in Tris-buffered saline. The following antibodies have been used: anti-GFAP (1:1000, chicken, ab4674, Abcam), Ki67 (1:500, rabbit, ab16667, Abcam), SOX2 (1:500, rabbit, PA1-094, Thermo Fischer Scientific Inc), Vimentin (1:100, chicken, ab24525, Abcam), NeuN (1:700, rabbit, ab177487, Abcam), connexin 43 (1:300, rabbit, 3512, Cell Signaling) and Nestin (1:200, rabbit, ab221660, Abcam). For staining with Nestin, antigen retrieval was performed by microwave for 10 min in 80° citric acid (0.1M citric acid, 0.1 M tripotassium citrate-2-hydrate in ddH_2_O). For all stainings, slices were incubated with the appropriate secondary antibodies and 4’, 6-diamidino-2-phenylindole (DAPI, 1:1000, ab228549, Abcam) for nuclear counterstaining.

Confocal microscopy:

samples were scanned with Zeiss confocal microscopes (CLSM 700, CLSM 800 and CLSM 900) equipped with ZEN 2011 (black edition, V7.1) using 10x air objectives (Plan-Apochromat 10x/0.45 M27, Zeiss), 25x oil immersed objectives (LCI Plan-Neofluar 25x/0.8 Imm Corr DIC M27, Zeiss) or 40x oil immersed objectives (Plan-Apochromat 40x/1.4 Oil DIC M27, Zeiss). For GFAP, Vimentin, Nestin, and SOX2 quantification, intensity in z-projected 30 μm stacks was measured in an area encompassing perilesional astrocytes after subtraction of background intensity in an adjacent unaffected region. The same area size and distance from the lesion core was used for the control condition.

### EdU administration and imaging

EdU was administrated via drinking water (0.4 mg/ml) immediately after antibody injection. Additionally, a single i.p. injection of EdU (5 mg/ml, 25 mg/kg BW) in NaCl was given. EdU-supplied water was provided for 7 days and then substituted for regular drinking water. Animals were perfused 14 days after lesion induction. EdU-supplied water was renewed at the latest after 3 days. Detection of EdU incorporation was performed using a commercially available detection kit (BCK-EdU-647, Click-iT, BaseClick). After the detection protocol, either additional stainings were performed as described above or DAPI alone in Tris-Triton buffer was added and left to incubate for 20 min.

#### Immunohistochemistry of human biopsy and autopsy tissue samples and image acquisition

Paraformaldehyde-fixed paraffin-embedded human CNS material was obtained from the archive of the Department of Neuropathology, University Medical Center Göttingen. A positive AQP4-antibody status was confirmed for the patients diagnosed after 2005; the status for one patient autopsied in 1962 is unknown. For this patient only clinical data are available, which indicate that the patient presented with acute myelitis and area postrema syndrome, both key features of AQP4-IgG-positive NMOSD. The diagnosis was histologically and immunohistochemically confirmed by the observation of lesions with damaged astrocytes, loss of GFAP and AQP4 as well as variable demyelination and macrophage infiltration. For our present study, 5-6 µm-thick slices were cut. Nuclear structures were labelled using hematoxylin. Immunohistochemical stainings were performed against GFAP (1:50, Dako M0761) and AQP4 (1:200, Sigma-Aldrich A5971). Proliferating cells were nuclear labeled with the Ki67 antibody (1:200, Dako GA626). The primary antibody binding was visualized using a biotinylated secondary antibody followed by developing with avidin-peroxidase and diaminobenzidine or 3-amino-9-ethylcarbazole.

Whole slide scans of GFAP-stained tissue slices were acquired at 200x-magnification using an Olympus VS120 slide scanner for quantification of multinucleated astrocytes. In order to reveal morphological features of single cells, high magnification (400x, 1000x) images of selected regions were acquired using an Olympus BX63 microscope.

### Correlative light and electron microscopy

#### Section preparation

Mice were perfused with a fixative containing 2% formaldehyde and 2.5% glutaraldehyde in phosphate buffer (0.1 M pH: 7.4) following the final imaging session. Brains were removed from the skull and postfixed in the same fixation solution for three hours, washed briefly in phosphate buffer and 100 µm-thick sections were collected with a vibratome (Leica VT1200 S). Sections were stained with nuclear staining DAPI (1:500) for 20 min, followed by imaging with a multi-photon microscope (Leica SP8 MP Dive Falcon), equipped with A 25x NA 1.00 water objective (HC IRAPO L 25x/1.0 W motCORR, Leica Microsystems). We acquired three fluorescence channels (excitation laser at 720 nm DAPI; excitation laser at 950nm eGFP; excitation laser at 1040 nm tdTomato) and used a transmission PMT to obtain additional structural information (location of blood vessels). Overview images of the whole sections were collected as well as complete Z stacks of ROIs (voxel size: 170 nm x 170 nm x 800 nm). The sections were placed back into fixative solution before further processing for electron microscopy.

#### Serial section electron microscopy

the vibratome sections were fixed with 1% OsO_4_ for 1 hour in 0.1 M cacodylate buffer at 0 °C, and in 1% aqueous uranyl acetate for 1 h at 4 °C. Samples were dehydrated in an ethanol series and embedded in Araldite/Epon (Sigma-Aldrich) 66% in propylene oxide overnight, 100% for 1 h at RT and polymerized at 60 °C for 20 h. The embedded samples were correlated with the transmission light microscope images acquired with the multi-photon images, to trim the ROI before the serial sectioning. Ultrathin serial sections (100 nm) were collected on silicon wafers using an ultramicrotome (Artos 3D, Leica Microsystems) equipped with a silicon wafer holder (CD-FH, Germany^63^). Sections were imaged in an Apreo 2 VS scanning electron microscope using the MAPS software package for automatic serial section recognition and image acquisition^64^ (Thermo Fisher Scientific). The following parameters were used for imaging: OptiPlan mode, T1 detector; pixel size of 4 nm or 7 nm, pixel dwell time of 3 or 5 µs, an electron high tension of 1.8 keV and a beam current of 0.1 nA. Serial section tiff images were aligned and the pairs of cells of interest were traced using the FIJI^57^-plugin TrakEM2.^65^ Traces and aligned images were imported for 3D reconstruction in Imaris (Oxford Instruments).

### Statistical analysis

Statistical analysis was performed in GraphPad Prism 9 and 10 software. Data is presented as mean ± SEM. Statistical significance was calculated using the Mann-Whitney test for comparing two unpaired groups and non-parametric ANOVA followed by Kruskal-Wallis test for comparing more than two groups. Statistical comparison of two paired groups was calculated by a nonparametric Wilcoxon signed-rank test. Comparisons of more than two paired groups were performed using a nonparametric Friedman test followed by Dunn’s multiple comparisons test. Correlation analysis was performed with a nonparametric Spearman’s correlation. P-values < 0.05 were considered statistically significant and indicated with asterisks in graphs as: * p < 0.05, ** p < 0.01, *** p < 0.001, and p *** < 0.0001.

### Materials and reagents

An extensive list of materials and reagents is provided in Supplementary Table 1.

## Data availability

Authors confirm that all relevant data are included in the paper/or its supplementary information files. Further information and requests for resources should be directed to the lead contact, Bruno Weber. This study did not generate new unique reagents. Mouse lines can be requested from the providing investigators and are protected by standard MTAs. Data related to human tissue analysis is available from CS upon request. This paper reports an original Python code for vector-based morphological analysis that has been deposited at the Mendeley repository (doi: 10.17632/xw8fv8gt8f.1). Source data are provided with this paper.

## Acknowledgments

We thank S. Weber, H. Osswald, N. Binini, and A. Siebert for technical and administrative support. We thank Jean-Charles Paterna and the Viral Vector Facility of the Neuroscience Center Zurich (ZNZ) for viral productions. We thank N. Snaidero for comments on EM data. We thank Ursula Lüthi for technical support with the CLEM experiments. The Aldh1l1^GFP^ mouse strain [STOCK Tg(Aldh1l1-EGFP)OFC789Gsat/Mmucd; identification no.: 011015-UCD] was obtained from the Mutant Mouse Regional Resource Center, a NCRR-NIH-funded strain repository, and was donated to the MMRRC by the NINDS funded GENSAT BAC transgenic project. MH was supported by the Deutsche Forschungsgemeinschaft (DFG) research grant (#444138499) and DFG transregional collaborative research center TRR 274/1, Project ID 408885537 B03, “Checkpoints of CNS recovery”, by the UZH Candoc Postdoc Grant (#FK-22-048) and the Swiss National Science Foundation (SNSF) Ambizione Grant (PZ00’3_216616/1). MTW was supported by the Stiftung für wissenschaftliche Forschung UZH (STWF-22-013). CT held a clinician scientist position of EXC 2067/1-390729940. ASS was supported by the SNSF (Eccellenza 187000). JLB was supported by the National Eye Institute (R01EY022936). S.J. was supported by the SNSF (TMAG-3_209272 / 310030_196869). CS was supported by the DFG transregional collaborative research center TRR 274/1, Project ID 408885537 B03, “Checkpoints of CNS recovery”, STA 1389/2-1, STA 1389/5-1 and the DFG under Germany’s Excellence Strategy (EXC 2067/1-390729940). TM was funded by the DZNE and by the DFG via EXC 2145 (SyNergy, ID 390857198), Mi 694/9-1 (Immunostroke, ID 428663564) and TRR 274/1,2 (B03, ID 408885537). BW was supported by the SNSF (31003A_156965).

## Author contributions

MH, MTW, TM and BW are responsible for the concept and study design. MH, MTW, NBS, JC, LR, JMMM, AK, GB, CT, CS, JLB, ASS, SJ were involved in sample/data acquisition and analysis. MH, MTW and BW drafted the manuscript and figures with input from all the authors.

## Additional information

The online version contains Supplementary Information material.

**Correspondence and requests for materials** should be addressed to the lead contact, Bruno Weber.

## Supplementary information

**Movie 1. Perilesional astrocytes show a transient calcium hyperactivity**

Time lapses of GCAMP6s of a perilesional reactive astrocyte (shown in Fig. 2h) at day 5 after AQP4-IgG injection (left panel) compared to astrocytes from a Ctrl-IgG-injected area (right panel); time in min:sec format (scale bar = 20 µm).

**Movie 2. Stack of a multinucleated astrocyte in vivo**

Animated z-stack (104 µm) of a perilesional reactive astrocyte 30 dpi (4-cell clone), shown in Fig. 3d. TdTomato+ astrocyte (red) with multiple nuclei (yellow arrowheads). Nuclei stained with Hoechst 33342 (cyan) *in vivo*.

**Movie 3. Awake two-photon time lapses of a perilesional astrocyte**

The mature astrocyte from Fig. 4f (red arrowhead) becomes reactive and gives birth to two daughter cells: one stays in close proximity (yellow arrowhead) and the other migrates via nucleokinesis to a new territory (cyan arrowhead); time in hours (scale bar = 20 µm).

